# Deformation microscopy for dynamic intracellular and intranuclear mapping of mechanics with high spatiotemporal resolution

**DOI:** 10.1101/403261

**Authors:** Soham Ghosh, Benjamin Seelbinder, Jonathan T. Henderson, Ryan D. Watts, Alexander I. Veress, Corey P. Neu

## Abstract

Structural heterogeneity is a hallmark of living cells and nuclei that drives local mechanical properties and dynamic cellular responses, including adhesion, gene expression, and differentiation. However, robust quantification of intracellular or intranuclear mechanics are lacking from conventional methods. Here, we describe new development of *deformation microscopy* that leverages conventional imaging and an automated hyperelastic warping algorithm to investigate strain history, deformation dynamics, and changes in structural heterogeneity within the interior of cells and nuclei. Using deformation microscopy, we found that tensile loading modes dominated intranuclear architectural dynamics in cardiomyocytes *in vitro* or myocytes *in vivo*, which was compromised by disruption of LINC complex molecule nesprin-3 or Lamin A/C, respectively. We also found that cells cultured on stiff substrates or in hyperosmotic conditions displayed abnormal strain burden and asymmetries compared to controls at interchromatin regions where active translation was expected. Deformation microscopy represents a foundational approach toward intracellular elastography, with potential utility to provide new mechanistic and quantitative insights in diverse mechanobiological applications.

## Introduction

Mechanobiology is an emerging field that describes how mechanical forces modulate the morphological and structural fitness of tissues, with applications in development, homeostasis, disease, and engineering of biological systems (*1*). Critical biophysical parameters in mechanobiology, microscale deformations, arise from mechanical loadings that impact the structural heterogeneity characteristic of the intracellular and intranuclear architecture. The deformation at multiple (e.g., extracellular, cell, nucleus) levels can trigger unique biochemical pathways and mechanotransduction cascades, and regulate diverse processes including adhesion and differentiation (*2*).

In recent years, specific studies and interest have focused on nuclear mechanobiology (*3–5*). The cell nucleus is known to contain, maintain, and interpret the genomic information that forms the basis for the characteristics of every individual. This information, encoded in the form of DNA, is highly organized within the nucleus to enable fast and accurate responses to cell stimuli. While maintaining spatial organization, the nucleus is also subject to significant deformation through direct physical connections to the cytoskeleton, via LINC (linker of nucleo- and cytoskeleton) complexes (*6, 7*), and consequentially to the extracellular environment, especially in mechanically challenged tissues such as heart, skeletal muscle, and cartilage. Through these interconnections, mechanical cues at the tissue level can be transmitted to and actively processed within the nucleus to influence, e.g., gene expression via activation of transcription factors (*8*), cell differentiation (*9*), cancer cell migration through constricted spaces (*4*), nucleoskeleton rearrangement (*3, 10*), and reorganization of the chromatin architecture (*11, 12*). The mechanics of the nucleus is also intertwined with biological functions, including transport through the nuclear membrane (*13*), nuclear rheology (*14*) and nuclear stiffness (*15*). Despite growing interest in nuclear mechanobiology, conventional methods to quantify deformation and mechanics do not highlight spatial mapping which could otherwise help elucidate regulation patterns of gene expression with respect to genomic organization and activation.

Numerous important methods have been described to measure cell mechanics, each with their own strengths and limitations (*16*). For the nucleus, the most commonly used techniques are based on the quantification of simple morphological changes such as aspect ratio, volume, or a characteristic dimension (*17, 18*). This type of analysis only considers geometric changes of the nuclear periphery and does not provide any intranuclear spatial information. Edge detection, fluorescence anisotropy (*19*), and texture correlation (*18, 20*) techniques provide only low resolution, spatial information and are typically limited to two dimensions. Moreover, they are overly sensitive to noise, and often require sharp spatial features that are typically not well defined in image data. In contrast, deformable image registration offers a more rigorous approach as it can quantify intranuclear deformation at high spatial resolution as demonstrated in intact tissue systems (*20*). Here, local differences in image intensities are used to match a discretized template image with a (deformed) target image to obtain deformation maps required for complete registration (*21*). Over the course of the analysis, a penalty factor (*21*) is used to enforce the registration. Additionally, regularization can be achieved using a hyperelastic material model. If known, the associated material properties can be assigned to improve the strain estimates in regions where image intensity differences may be lacking. Spatial averaging in the form of normal or Gaussian blurring are used in the image analysis to avoid local minima which would stop the global image registration prematurely resulting in false, unreliable deformation data (*22*). Consequently, defining a suitable method to map deformation, or a correct set of parameters to obtain an optimal deformation map in a manageable time frame, is challenging and largely lacking.

We introduce a technique, *deformation microscopy*, which utilizes an automated sweep over a wide range of registration parameters, to quantify precise, high-resolution, and reliable spatial patterns of intracellular displacements and strain. We demonstrated and validated the efficacy of the technique across several biological scales, including examples of extracellular matrix, cell, and nucleus deformation *in vitro* and *in vivo*. Next, we focused on applying the technique to understand the spatiotemporal mechanics of nucleus in several normal and pathological conditions. We altered the integrity of nuclear envelope by modulating the KASH domain, nesprin-3, and Lamin A/C to understand their structural role in nuclear mechanics both *in vitro* (cultured cells) and *in vivo* (living mice). Further, we revealed the compromised nuclear mechanics in cardiomyocytes cultured in fibrotic, pathological conditions. We additional showed how new deformation microscopy can be combined with independent measures like traction force microscopy, to reveal new biological insight into cellular stain burden and asymmetry during hyperosmotic loading.

## Results

### Deformation microscopy is a versatile tool to quantify intracellular and intranuclear biomechanics

*Deformation microscopy* is based on the minimization of an energy functional *E*(*X, φ*), a function of space *X* and deformation map *φ* which is the difference between the deformation induced energy *W* and the image comparison based energy *U* (Fig. 1a and Supplementary note 1). In a basic implementation, *z*-projections of template (undeformed) and target (deformed) images are used to quantify local deformations (Fig. 1a, Supplementary Fig. 1). Images are meshed and the hyperelastic warping algorithm is applied to find the deformation map. A sequential spatial filter is applied to overcome the local image registration-driven premature registration (Supplementary note 2.2). The deformation map can be used to generate various strain measures in the object of interest as suitable for the specific application, here demonstrated using cell nucleus as an example (Supplementary note 3). The material stiffness of the nucleus and the penalty factor(*λ*) are the two key parameters that are changed to find the best deformation map through optimization of root mean square error (RMSE) of image pixel intensity derived from template and target images (Supplementary note 2.1 and 2.3, Supplementary Fig. 2). A custom-built code automated the process to measure intranuclear deformation by using widely-available computational resources. One set of calculations averaged 5-30 minutes, depending on the complexity of the specific application. We quantified the deformation maps of a single beating cardiomyocyte *in vitro* (Fig. 1b) and in skeletal muscle tissue *in vivo* using image information before and after contraction, respectively, to demonstrate the utilization of the deformation microscopy algorithm across different scales and applications (Fig. 1c). The accuracy and reliability of the technique was validated for all image types but here, we demonstrated the validation specifically for the nucleus over a wide range of possible material stiffness assigned to heterochromatin and euchromatin spaces (Fig. 1d) by thresholding the image with a decided cut-off intensity value. The forward (known) deformation model applied a uniform normal force on the nuclear periphery to simulate a deformation map. As shown (Fig. 1e), *deformation microscopy* was capable of reliable strain and chromatin organization quantification over an order of magnitude range of material property values. Euchromatin (E) is thought to have lower stiffness than heterochromatin (H), and hence the comparatively less accurate registration was observed for a low material property ratio (i.e. H:E = 1:10) case where some elements near the nuclear periphery had overestimated strain values. This was further quantitatively validated by comparing known deformation with computed deformation (Fig. 1f), where Nucleus B (H:E = 10:1) yielded the most accurate measurement followed by Nucleus C (H:E = 2:1). While this technique provided detailed intranuclear deformation maps, bulk mechanical measurements alone result in unreliable strain quantification (Supplementary Fig. 3.a). Further, the deformation in intranuclear space was found be related to the local chromatin density, and hence to the image intensity (Supplementary Fig. 3.b-d). The sensitivity of the deformation measurement with regard to the key parameters material stiffness and penalty factor was also analyzed (Supplementary Fig. 4), and we found that, based on the specific application, an optimal combination of those two parameters lead to the best registration results. From the high-resolution principal strain vector map (Fig. 1.a), it was evident that this method used the small image features for the strain measurements and was limited only by the image quality obtained by the available imaging modality (e.g. microscopy).

**Figure 1:**
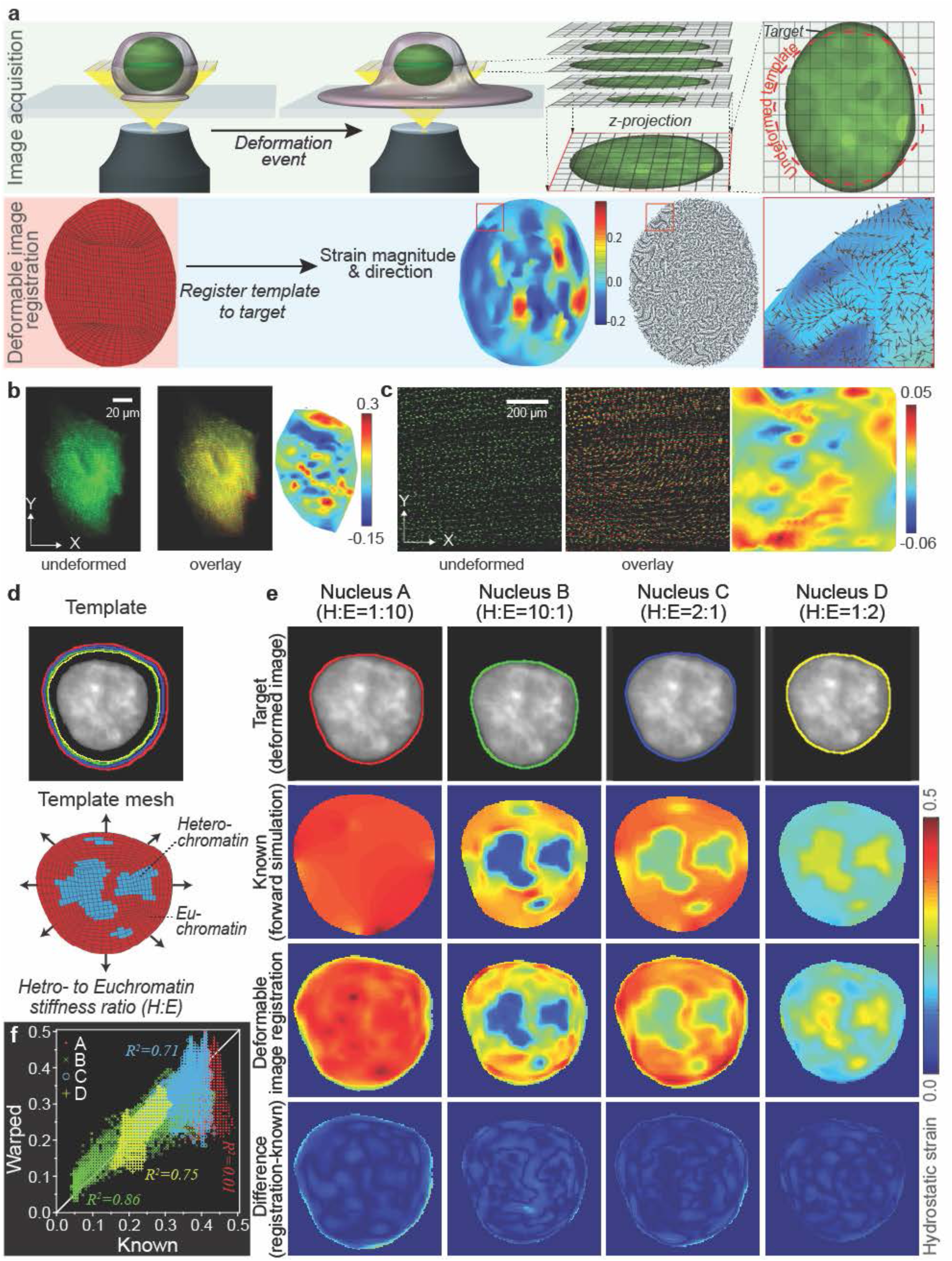
Strain measurement using *deformation microscopy* at multiple (tissue/matrix to subcellular) biological scales and its validation. (a) Illustrated deformation microscopy workflow for generating strain maps, specifically elaborated in nucleus. Nuclear *z*-stacks in undeformed (template) and deformed (target) state are acquired using confocal microscopy. *Z*-projections are then generated to carry out the deformation quantification in a computationally less demanding two dimensional framework. After a mesh for the template nucleus is generated, a mathematical model uses the iterative hyperelastic warping algorithm to quantify intranuclear strain by registering the undeformed image to the deformed image. The result is a high resolution spatial deformation map. Shown here is a deformation surface map and deformation vector directions. Overlap of deformation value and direction is shown in the zoomed in panel for the red inset area. (b) Embryonic cardiomyocytes where transduced with lentiviral particles harboring an α-Actinin-mRuby2 construct under the control of an EF1α-promoter. The labeled sarcomeric aparatus of a cell was then imaged during spontaneous contractions and the image stack was used to calculate strain maps via deformation microscopy. Undeformed (green) and overlay (undeformed: green + deformed: red) cardiomyocyte is shown. Strain map shows the E_yy_. (c) Skeletal muscle was stained with NucBlue live DNA stain and contracted through electro-stimulation *in vivo*. Stained nuclei were used a visual nodes to demonstrate the feasability of the technique on the tissue scale. Undeformed (green) and overlay (undeformed: green + deformed: red) tissue is shown. Strain map shows the hydrostatic strain. (d) Validation of the deformable image registration technique specifically elaborated in nucleus. The template nucleus was subjected to uniform outward normal force along the nucleus perimeter. The intranuclear region was meshed and segmented into a low density euchromatin region (red area) and a high density heterochromatin area (blue area). Green, yellow, red and blue outlines show the deformed target nucleus perimeter for different heterochromatin to euchromatin stiffness ratios (H:E). (e) Top row shows the deformed template images. Hydrostatic strain maps are shown using forward finite element simulation (second row) or using deformable image registration (third row). The bottom row shows the difference between forward simulation and deformable image registration. Note that the difference was near zero throughout the nuclear domain. (f) Quantitative comparison of hydrostatic strain maps generated by deformable image registration compared to known strains from forward simulations. The best match is obtained for phyiologically relevant heterochromatin to euchromatin stiffness ratio of 2:1 (R^2^ = 0.86).

### Role of intranuclear-specific tensile loading modes, and LINC complex KASH domain proteins and nesprin-3, is revealed during cardiomyocyte beating *in vitro*

We showed that the LINC complex proteins at the nuclear periphery played a crucial role in the strain transfer from cell to nuclear space, and in regulating key tensile loading in active force-generating cells. We conducted intranuclear strain measurements in the context of cyclic loading of cardiomyocyte (CM) nuclei during cardiac contractions (Fig. 2). The CMs were derived from embryonic mice (E16.5) that harbored an eGFP-tag on the H2b histone (*H2b-eGFP*) (see online Methods for details). Embryo-derived CMs have the advantage of developing spontaneous beating behavior within 1 day of seeding and culture.

**Figure 2:**
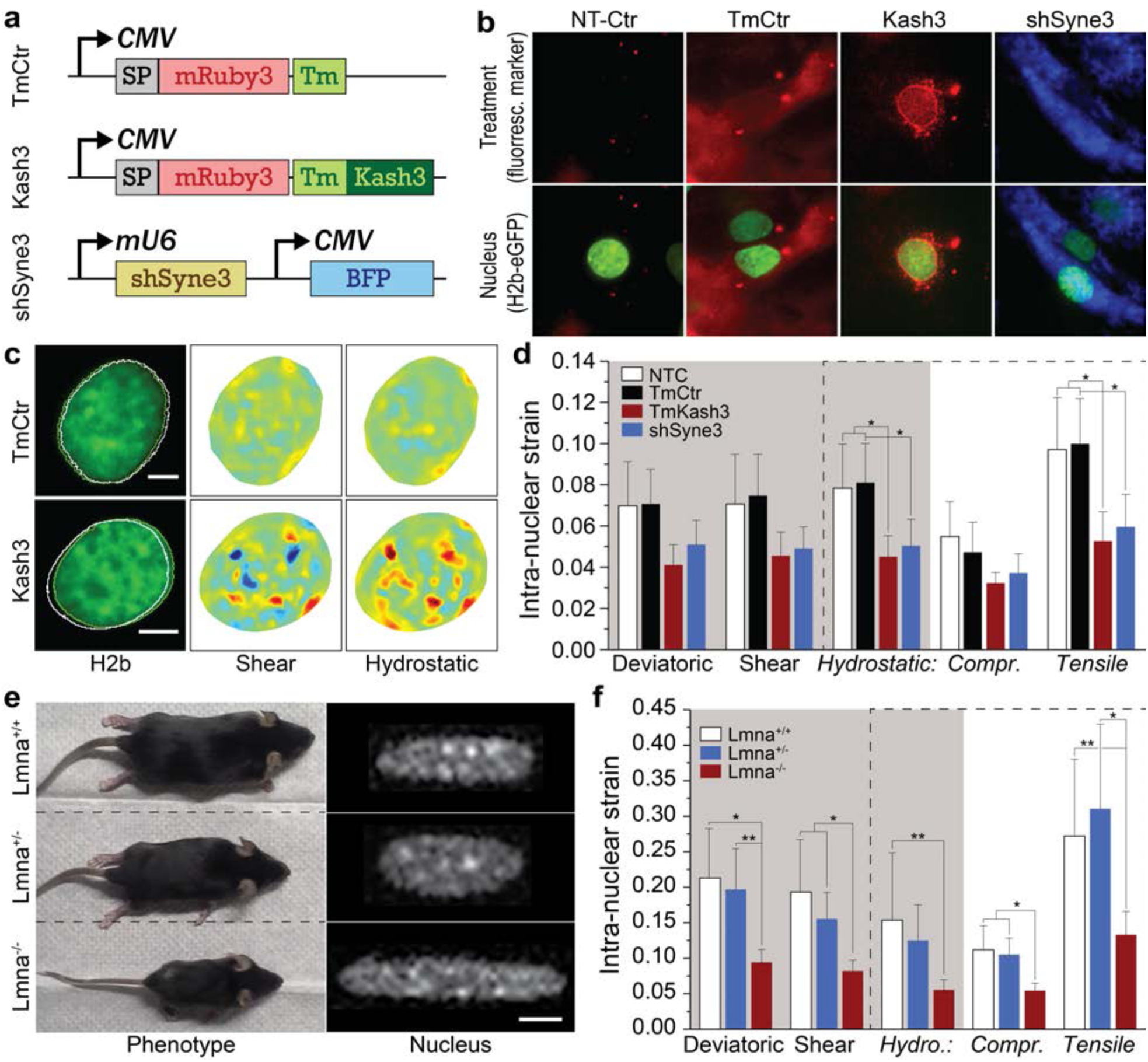
Disruption of LINC complex molecule nesprin-3 in cardiomyocytes *in vitro*, and knockout of Lamin A/C in myocytes *in vivo*, reveals altered strain transfer to the nucleus. (a) Lentiviral constructs designed to disrupt LINC connections; SP=signal peptide, Tm=transmembrane domain (b) Primary CMs from H2b-eGFP expressing embryonic mice 4 days after transduction; NT-Ctr=non-transduction control. (c) Nuclear strain maps from *deformation microscopy* analysis for a decoupled (sR3-Kash3) and a control nucleus (sR3-TmCtr); white lines indicate nuclear boundary at peak contraction. (d) Differential strain analysis of decoupled and control CM nuclei; n=6; error bars=standard deviation; (* p < 0.05). (e) Overall physiological and nuclear phenotype of hemi- and homozygote *LMNA* deficient mice compared to wild type littermate. (f) *In vivo*, for same tissue strain (pre-calibrated), nuclear strain in skeletal muscle is significantly lower in *LMNA*^*-/-*^ mice compared to *LMNA*^*+/+*^ and *LMNA*^*+/-*^ mice; scale bar=5 µm; n=9. (* p < 0.05, **p < 0.01).

We were interested to understand how intranuclear strain would change after disrupting elements associated with the nuclear membrane. LINC (<u>Li</u>nker of <u>N</u>ucleoskeleton and <u>C</u>ytoskeleton) complexes are broadly known to play a critical role in strain transfer from the cell periphery into the nucleus as they connect the cytoskeleton to the nucleoskeleton. LINC complexes are comprised of SUN and nesprin proteins: SUNs bind the nuclear lamina inside the nucleus, stretch through the inner nuclear membrane and connect to nesprins in the perinuclear space via highly conserved KASH domains (*7*). Nesprins in turn span through the outer nuclear membrane and connect to different parts of the cytoskeleton. In CMs, intermediary filaments are mainly represented by desmin which encases all sarcomeres in a honeycomb like fashion and laterally integrates with costameres (*23*). Nesprin-3 was specifically targeted here as it connects to intermediary filaments.

We hypothesized that disrupting one or multiple LINC complexes should significantly alter intranuclear strain intensities in the previously introduced CMs model. For that, CMs were transduced with lentiviral particles harboring a dominant negative sR3-Kash3 construct (Fig. 2a, b), which disrupts the connection between the nesprin-3 and intermediate filaments. A control construct sR3-TmCtr was designed which was identical to sR3-Kash3 but lacked the KASH domain and hence was not be able to compete with nesprin-3 for SUN connections. Additionally, we designed the lentiviral vector shSyne3 which carried an shRNA expression cassette that specifically targeted nesprin-3 and a blue fluorescent protein (BFP) to identify transduced cells (Fig. 2 a, b; Supplementary Fig. 5). Transduction of CMs with sR3-Kash3 showed that it formed a distinct ring around the nuclear border as it replaced nesprins. In contrast, sR3-TmCtr formed only a faint ring around the nucleus and was otherwise distributed throughout the cell as it most likely diffused along the ER continuum away from the nucleus. Since KASH is highly preserved between species and isoforms, this dominant negative construct led to an unspecific disruption of all LINC complexes. Immunostaining further confirmed the absence of nesprins 1 and 2 around the nuclear border in sR3-Kash3 transduced, but not sR3-TmCtr transduced CMs (Supplementary Fig. 5).

To provide a comprehensive analysis using *deformation microscopy*, we generated and compared hydrostatic, shear and deviatoric strain maps for all experiments. Average of absolute strain was computed for all three strain types: (avg(abs(ε_hyd_))), (avg(abs(ε_dev_))), (avg(abs(ε_shear_))). For the hydrostatic strain, further classification was performed only with the tensile and compressive components: (avg(abs(ε_hyd, tensile_))), (avg(abs(ε_hyd, compr_))). Tensile strains were compromised with LINC complex disruption. Intranuclear strain fields were reduced across all strain types in sR3-Kash3 and to a lesser extend in shSyne3 transduced CMs compared to non-transduced CMs. In contrast, sR3-TmCtr transduced cells showed similar strain intensities compared to non-transduced CMs.

### Lamin A/C maintains tissue-to-nucleus strain transfer in skeletal muscle activation *in vivo*

Next, we showed that the lack of lamin A/C, a structural protein in the nucleus, leads to altered tissue-to-nucleus strain transfer *in vivo* in a murine model (Fig. 2). Lamin A/C, along with lamin B1/B2 creates a meshwork under the nuclear membrane. *In vivo* muscle active stimulation lead to a complex loading mode on individual nuclei (Fig. 2e-f). Lamin A/C is thought to provide a structural scaffold to the cell nucleus, with the SUN proteins and the chromatin are connected to the lamin A/C associated proteins in a complex manner, still to be understood. *In vitro*, strain transfer experiments suggest a higher strain transfer to lamin A/C deficient nuclei, and thus a lower stiffness of nuclei (*15*). On the contrary, AFM studies on single cell nuclei suggested a stiffer nucleus in lamin A/C deficient cells (*24*). Therefore, the mechanical role of lamin A/C in the nucleus, especially *in vivo* is far from settled. We found that for the same tissue deformation, all the measures of nuclear strain decrease with lack of lamin A/C. Lamin A/C deficient (Lmna^-/-^) nuclei show significantly lower strain than the Lmna^+/+^ and Lmna^+/-^ counterparts. For all strain measures, the average absolute strain dropped by 40-50% in Lmna^-/-^ with respect to Lmna^+/+^. When the tensile and compressive hydrostatic strain was separated and compared among the groups, they did not show much preferential difference, because of a complex loading mode on the nuclei *in vivo*. Interestingly, the Lmna^+/+^ and Lmna^+/-^ groups did not show any significant difference in nuclear strain. This finding supports the observation that Lmna^+/+^ and Lmna^+/-^ mice do not exhibit any significant difference in development, phenotype, and viability.

### Fibrotic cardiomyocytes display abnormal strain burden at interchromatin regions

Using a normal and pathological model of cardiomyocyte we showed that the strain burden taken by distinct interchromatin regions is compromised in fibrotic nucleus (Fig. 3). It is known that CMs modulate their beating behavior based on the stiffness of their environment with optimal beating properties being observed on substrates that resemble the healthy native heart stiffness (10-20 kPa) whereas contractile forces decline on stiffer substrates that mimic pathologically altered tissues, e.g. due to fibrosis or after a myocardial infarction (*25*). This decline in beating activity is argued to be a direct result from altered nuclear mechanosensation triggered by the increased substrate stiffness (*25*), however, no investigation of cardiac intranuclear strains and their change with altered mechanical environment as seen in disease has been conducted so far. We generated intranuclear strain maps of *H2b-eGFP* CMs that were plated on silicon substrates with either soft (∼15 kPa) or stiff (∼400 kPa) elastic properties to mimic native and pathological environments. The histone tag allowed for a contrast-rich live-imaging of the nucleus during CM contraction. As H2b is present in all nucleosome complexes, it can be used as a direct marker for local chromatin compaction. Four days after plating, image series with ∼6.4 fps were acquired over a period of 10s during which photobleaching was negligible. For each nucleus, one contraction cycle was picked in which to perform in-depth intranuclear strain analysis. Image frames in the post-diastolic resting states were selected as undeformed reference images (template) and t_1_-t_4_ represent the deformed states during the contraction cycle. In CMs cultured on a stiff substrate, CM contraction cycles were shortened as expected, hence image series typically resulted in only three deformed images during one contraction cycle, as compared to four in the soft case.

**Figure 3:**
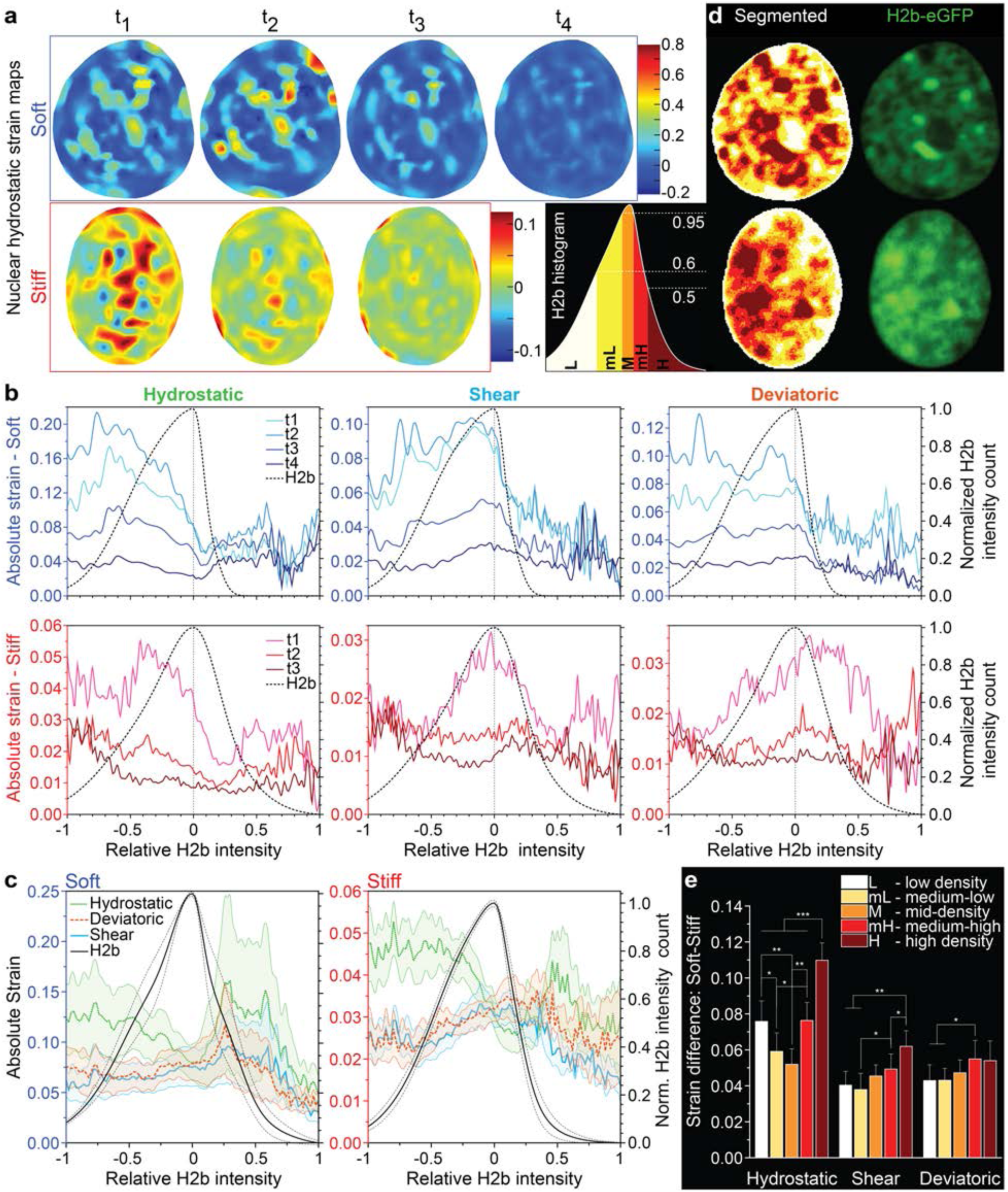
Interchromatin regions display altered strain burden in cardiomyocytes under pathological culture conditions. Comparative assessment of intranuclear strains in cardiomyocytes (CM) plated on soft (normal) versus stiff (pathological) substrates. Primary CMs from embryonic mice expressing *H2b-eGFP* were plated on either soft (∼15 kPa) or stiff (∼150 kPa) PDMS substrates and image stacks of nuclear deformation during CM contraction were recorded. (a) Hydrostatic strain maps were generated via *deformation microscopy* using fluorescence image stacks with diastolic resting state as undeformed template. (b) Strain dynamics along normalized chromatin compaction. Strain maps were spatially correlated with *H2b*-chromatin intensity maps and strains of pixels with similar intensity where averaged and plotted over this intensity. To allow for comparison, the chromatin intensity peak of each distribution was set to 0 and both sides were linearly normalized to −1 (lowest value) and +1 (highest value). Different lines present different time steps as seen in (a). (c) Averaged absolute nuclear strains at peak contraction over chromatin compaction for n=5 cells (each); areas indicate S.E.M. (d) Left lower panel: Representative segmentation of chromatin intensity histogram into low (L), mid-low (mL), mid-density (m), mid-high (mH) and high (H) density areas. Mid and right panel: resulting segmentation map and corresponding nuclear *H2b-eGFP* intensity map. (e) Difference in absolute strain of CMs plated on soft or stiff substrates for each density segment. Error bars present STD; *** p<0.001; ** p<0.01; * p<0.05.

Nuclear hydrostatic strain maps derived from a CM nucleus plated on soft substrate displayed high tensile and compressive regions during systolic peak contraction (t_2_ for soft, t_1_ for stiff), most likely due to combined tensile forces exerted from the cytoskeleton and overall compressive forces due to cell shortening (Fig. 3a-c, supplementary video 2). The overall strain burden then gradually decreased during diastolic relaxation. For the nuclei plated on the stiff substrate, strains followed a similar trend, however, with amplitudes that were almost one order of magnitude lower compared to the soft substrates due to the previously mentioned stiffness-induced decline of CM contraction forces. With both high tensile and compressive strains, this model was well suited to demonstrate that bulk strain analysis methods highly underestimate actual strain values because they allow local intranuclear strains to cancel each other out by only considering deformation at the nuclear boundary. To support this idea, we compared averaged absolute hydrostatic strains generated from our *deformation microscopy* analysis (avg(abs(ε_hyd_))) with bulk strains calculated from the average change in minor and major axis (abs(ε_bulk_)) utilized by previous researchers(*8*) (Supplementary Fig. 6). To further show that the underestimations in bulk strain are a result of neglecting intranuclear strains, we calculated another value were hydrostatic strains were first averaged and then made absolute (abs(avg(ε_hyd_))) to allow for the same effect. As expected, averaged absolute local strains were twice as high compared to bulk strains for both soft and stiff (soft: 0.111 vs 0.046 and stiff: 0.032 vs 0.018) while hydrostatic strains that where first averaged then made absolute showed almost identical values. It is important to mention that these local extrema can be up to 10× higher compared to averaged hydrostatic strains or 20× higher compared to bulk strain estimated, respectively.

Besides providing more reliable average strain data, spatial strain maps can provide new detail and insight into nuclear mechanics through combined analysis with other spatial information. Here, we utilized the fluorescence intensity of the *H2b* histone tags, to perform a differential analysis of strain types along the chromatin compaction continuum. Since fluorescence intensity values are arbitrary, we generated chromatin intensity histograms that result in a bell curve with a prominent peak that can be fitted onto a 3-term Gaussian distribution. Chromatin intensity values of the histograms were then rescaled to [-1 +1] with −1 being the lowest density, +1 the highest density, and the histogram peak set to 0 to compare strain data from different nuclei. As shown in Fig. 3b, chromatin intensity correlated strain data for one set of CM nuclei plated on either soft or stiff substrates over time. We observed that different strains varied in prominence along different chromatin compaction states during a contraction cycle. For example, changes in hydrostatic strains were more significant in lower density regions, but became less significant towards the mid-density regions around the histogram peak and stayed low in high-density areas. In contrast, most prominent shear and deviatoric strains changes coincided more with low- and mid-density chromatin, and were low again for high-density areas. These contrasting strain trends with hydrostatic strains being low, and shear and deviatoric being high, around mid-density chromatin areas was further confirmed in the analysis of *n*=5 CM nuclei for each substrate (Fig. 3c and Supplementary Fig. 6). While a similar trend was observed for CMs plated on stiff substrates (with lower amplitudes) closer segmentation analysis revealed significant differences between density bins. To allow direct comparison of different chromatin density regions, chromatin intensity histograms were binned into 5 areas with normalized intensity intervals between [0; 0.6] for low-density, [0.6; 0.95] for medium low-density, [0.95; 0.95] for mid-density, [0.95; 0.5] for medium high-density, and [0.5; 0] for high-density chromatin (Fig. 3d). Intervals where chosen to provide roughly equal numbers of data points for each density region. Binned chromatin intensity maps of both nuclei showed a distinct pattern of low-density chromatin accumulating at the nuclear border and dense chromatin cores distributed throughout the inner nuclear area with medium-low and medium-high density chromatin suspended in between. Analysis of the difference in strain between nuclei of CMs plated on soft versus stiff substrates for each density segment revealed most significant discrepancies for hydrostatic strains in mid density areas showing in average 5.2% lower strain for stiff compared to soft in mid-density areas while being double as low with an 11.0% decrease in high density areas. Less distinct discrepancies between density bins could be observed for shear and deviatoric strain values; however, stiff nuclei experienced significantly less strain in higher density bins compared to lower density bins.

### Organized stress fiber-rich primary apex creates strain asymmetry in cells during hyperosmotic loading

Finally, we showed that the primary apex generated an assymetric strain distribution in the nuclei of fibrotic cells, using chondrocytes as a model system for hyperosmotic loading (Fig. 4). We also demonstrated how the combination of two independent measurements in the intranuclear space (deformation microscopy) and extracellular space (traction force microscopy) can reveal novel mechanistic insight of passage 4 (P4) chondrocytes during hyperosmotic conditions (Fig. 4, Supplementary Fig. 7). Cartilage goes through cycles of hyperosmotic and hypoosmotic shock due to its poroelastic nature, and changes in osmotic pressure leads to intracellular water transport cell and nuclear deformation, and altered gene expression (*26*). To analyze spatially-dependent nuclear strain maps in this process, we imaged the progressive deformation of P4 chondrocyte nuclei for 12 min after hyperosmotic loading. We observed increasingly compressive nuclear strains with temporal progressions of hyperosmotic loading as determined by bulk mechanical strain analysis (Supplementary Fig. 7) due to the extrusion of water as mentioned above. To combine spatial strain distributions inside the nucleus with forces exerted on the cells substrate, chondrocytes were cultured on fluorescent bead-embedded PDMS. After 12 min of osmotic loading, cells were released from the substrate via trypsin to capture the undeformed state of the silicon substrate (Fig. 4a, Supplementary video 1). Traction force microscopy (*27*) was then used to calculate substrate traction force from bead displacements during hyperosmotic loading using the post-trypsin state as reference. Substrate traction forces declined, most likely due to cell shrinkage, as decreased cell area was previously correlated to loss of traction (*28*). In our studies, higher magnitudes of traction forces was observed near the three distinct cell apexes. Substrate stress relaxation maps could then be calculated from the difference in traction force using the pre-osmotic loading condition (0 min) as reference (Fig. 4b). Nuclear strains (Fig. 4c) revealed that shape change-related deviatoric strains were unrelated (R^2^ = 0.12) with substrate stress relaxation while shear (R^2^ = 0.97) and hydrostatic strain (R^2^ = 0.99) related at different time points. Note, however, that hydrostatic strain was three times greater in magnitude than the shear strain (Fig. 4d). This trend was verified by comparing strains at the loading end point (11 min) of three different cells.

**Figure 4:**
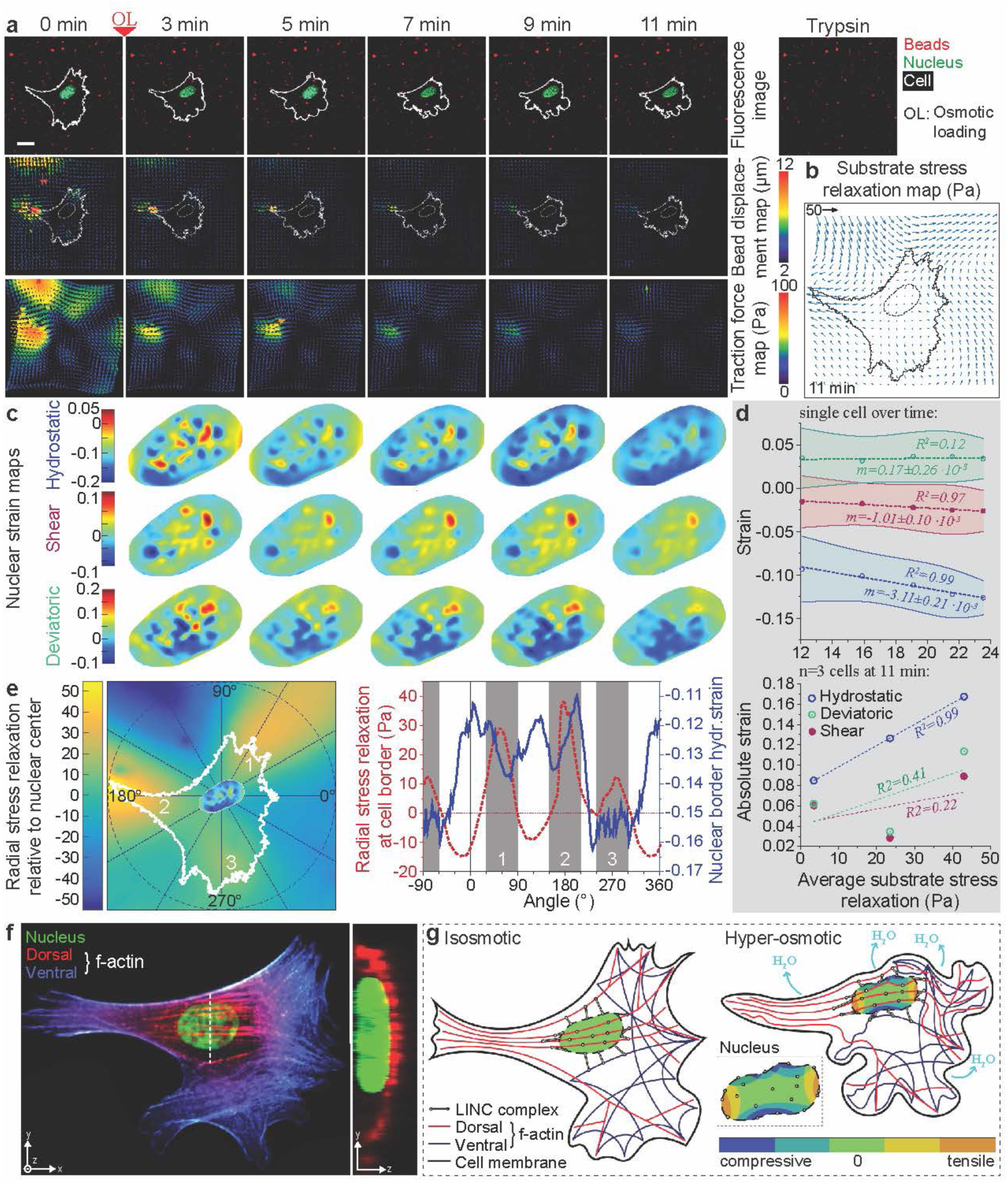
Combined quantitative deformation microscopy and traction force microscopy modalities at intranuclear and extracellular spaces reveal the role of primary cell apices in creating strain assymetry in the hyperosmotically-challenged chondrocyte nucleus. (a) Cells (white boundary) and nuclei (green) cultured on PDMS with embedded polystyrene beads (red) shrink due to hyperosmotic loading. Scale bar: 20 µm. Bead displacement maps (second row) and traction force maps (third row) over time were quantified using a reference image at 16 min obtained after cells were released through trypsin-treatment (Trypsin). (b) Substrate stress relaxation map at 11 min was calculated using the traction force map before hyperosmotic loading (0 min) as a reference. (c) Spatiotemporal hydrostatic, shear and deviatoric strains in nuclei over time are heterogeneous and mostly compressive, with some tensile spots. (d) Upper graph: Correlation of average nuclear strains with average substrate stress relaxation at different time point during hyperosmotic loading. Lower graph: Correlation of nuclear strains with average substrate stress relaxation for 3 different cells at the end-point of hyperosmotic loading (11 min); areas represent STD. (e) Left side: Radial stress relaxation map has been calculated from (b) by recalculating magnitude with respect to the nuclear center; nucleus inlay shows the hydrostatic strain map at 11 min, white outline indicates cell boundary and white numbers represent cell apexes. Right side: Simultaneous plotting of radial substrate stress relaxation at the cell boundary and hydrostatic strain at the nuclear border vs. their radial position with respect to the nuclear center. (f) *Z*-stack confocal imaging of a stained P4 chondrocyte: Nucleus (DAPI, green) and actin fibers (phalloidin). Actin staining has been split into two regions depending on the *z*-location: lower *z* (ventral, blue) and higher *z* (ventral, red). White dotted line indicates the position of the *yz*-projection on the right. Note that the position of the dorsal stress fibers is above the nucleus. (g) Model of observed nuclear hydrostatic strain during hyperosmotic loading. Strain patterns in (e) indicate a difference in the strain pattern along the primary apex 2 vs. secondary apexes 1 and 3. Apex 2 associated dorsal stress fibers keep the nucleus aligned along the apex-axis resulting into lower compression while the collapse of ventral stress fibers might contribute to higher nuclear compression at the secondary apex locations.

Next, to demonstrate the benefits of computing spatial strain maps, we investigated whether a spatial relationship existed between substrate stress relaxation and intranuclear hydrostatic strains. For this purpose, we generated a radial stress relaxation map with values indicating the stress towards the nuclear center (Fig. 4e). The radial stress relaxation at the cell border was then plotted together with the hydrostatic strain at the nuclear border (10% of outer nuclear area) against the radial position in perspective to the nuclear center. Unsurprisingly, substrate stress relaxation revealed three prominent peaks that coincided with the radial position of each cell apex, respectively. However, hydrostatic strain at the nuclear border showed a different trend with a prominent peak (lower compression) at the radial position of apex 2 but higher compression at apex 1 and 3 (Fig. 4). Additionally, a peak was observed between apex 1 and 2, and more distinctly, between 1 and 3 at the opposing radial position of apex 2. *Z*-stack confocal imaging of an actin stained P4 chondrocytes with a similar cell geometry showed that the most prominent apex in a cell, like apex 2, is distinct from the others as it serves as focal point for dorsal stress fibers that stretch over the nucleus and connect to ventral stress fibers on the opposite side, as in apex 1 and 3 (Fig. 4f).

## Discussion

Deformation microscopy enables detailed, spatially-dependent investigations of the cell and nucleus interior in a broad range of physiological and disease applications. Development of deformation microscopy leverages conventional imaging and an automated hyperelastic warping algorithm to investigate strain history, deformation dynamics, and changes in structural heterogeneity within the interior of cells and nuclei. Deformation microscopy represents a foundational approach toward intracellular elastography, and can potentially provide quantitative insights in diverse mechanobiological applications. The technique can be applied to data from any image modality, and only requires sufficient ‘texture’ in images representing critical subcellular structures, e.g. actin stress fiber or chromatin architecture within cells. Reliable spatial registration map generation is only limited by imaging quality.

Deformation microscopy is automated and valid for both 2D and 3D imaging modalities (*20*). Accordingly, the derived parameters (in this case, *strain*) can be quantified in 2D as well as in 3D. The automated, unbiased optimization capability in the present technique adds much repeatability and makes the technique user-independent. Therefore, the present technique is a significant advancement over our previous work (*20*), which required an experienced user to provide judgment to set optimization parameters that would maximize accuracy of the results. In the present study, we also intentionally utilized only cell models grown in 2D environments in the assumption that strains in *z*-direction will be negligible for such cases. However, extension to 3D is easily possible at a cost of increased computational demand.

Our model assumes a hyperelastic material model. Hyperelasticity encompasses a large class of material models, including complex and nonlinear elasticity, and enables description of intracellular deformation for most applications. For the nucleus, very high strain occurs in a few scenarios such as migration of cancer cells through constrictions (*4*), which then recover after passing through the constriction, thus still showing hyperelastic and not plastic or permanent deformation behavior. For most applications of nuclear mechanics, the deformation is relatively low or recovers quickly over time. Viscoelastic or time-dependent behavior of the nucleus has also been investigated in isolated cells and nuclei. It was found from two independent studies that the nucleus does show a viscoelastic behavior at the verge of a suddenly applied deformation (*3, 29*). After a short stabilization period, the nucleus behaves as an elastic solid. Those studies induced sudden deformation on nuclei using micropipette aspiration or bead displacement. *In vivo*, sudden deformation on nuclei rarely happens and thus, an elastic material model can be appropriate. Importantly, because biological materials often also display time-dependent viscoelastic behavior, and plastic behavior at very large deformation, mechanical characterization of these materials is still possible via deformation microscopy through the acquisition of time course data.

Our technique provides mechanistic insight into the nuclear mechanics by LINC disruption and LMNA knockout study. Though interrupting the of KASH domain can alter the nuclear mechanics as expected because of its connectivity to all four nesprin types, the underexpression of only nesprin-3 leads to drastic consequences in nuclear mechanics. The mechanistic roles of nesprins in cell biology are still being discovered. Out of the four types of nesprins, nesprins 1 and 2 are mostly investigated in the context of nuclear mechanics and mechanobiology (*30*). The loss of nesprins 1 and 2 are known to be responsible for several diseases including Emery-Dreifuss muscular dystrophy and dilated cardiomyopathy. The loss of nesprin-4 is related to hearing loss. Interestingly, no disease has been reported so far where nesprin-3 is compromised, thus making it the least investigated. However, the ubiquitously conserved nature of nesprin-3 also shows how important nesprin-3 is to fundamental biological processes. In endothelial cells, nesprins are known to play an important role in the response to fluid shear stress, with depletion of nesprin-3 causing altered cell morphology and impaired cell polarization and migration in the direction of the fluid flow (*31*). Another work demonstrated that the actomyosin, nesprin-3, and vimentin cooperatively move the nucleus like a piston during 3D migration (*32*). Therefore, the role of nesprin-3 in nuclear mechanics is increasingly important, and we show for the first time how the lack of nesprin-3 compromises cardiomyocyte contractility.

The lack of lamin A/C is involved in numerous diseases, collectively known as laminopathies (*30*). Our study confirms the abnormal nuclear strain *in vivo* is similar to *in vitro* analyses. The compromise in lamin A/C is known to cause abnormal nuclear mechanics and phenotype (*33*). Besides just mechanically softening or stiffening of nucleus, lamin A/C may probably have a role in chromatin organization, homeostasis and epigenetic regulations, and overall gene expression. Thus, this study advances the technology to study the role of lamin A/*C in vivo*, which is more relevant from the clinical perspective, and directly relates loss of this structural molecule to intranuclear patterns of deformation.

We provided the first differential nuclear strain analysis for CMs in different mechanical environments to find support for the hypothesis that nuclear mechanosensation is linked to the well described beating decline of CMs on stiffer substrates. It has been shown that locally applied strains, particularly shear and hydrostatic strain, are sufficient to alter the expression of genes without any other influence (*5*). It is further known that the activity of a gene is related to its compaction state, which is orchestrated by epigenetic modifications (*34*). For example, methylations of different histone residues can inactivate genes that were important during development but are locked away in the adult stage. Cardiac performance decline in conditions such as hypertrophy, ischemia, hypoxia and atrophy have long been related to the reactivation of fetal gene networks (*35*). If we further consider that high density chromatin areas are mainly comprised of silenced heterochromatin, as dense compaction is one of the hallmarks of heterochromatin, we can postulate that the elevated differences in strain burden of high density areas, as observed in this study, could have a profound impact on epigenetic controls by disturbing the balance of silenced developmental genes and active genes subsequently leading to beating abrogation. These results form a strong basis to further investigate the effects of altered nuclear mechanics on chromatin organization (and subsequent overall control of the genome) not only in heart disease, but many other conditions that are marked by dramatic changes in mechanical properties of the respective tissues (e.g. diverse fibrotic conditions in the brain or liver, osteoarthritis in joint tissues, cancer).

Deformation microscopy, in combination with other established microscale measurement techniques, can provide interesting mechanical insights unavailable from the use of either technique alone. In our studies, we postulated a model where dorsal actin fibers keep the nucleus tensed along the primary apex axes while allowing for higher compression at the sides. Based on these experimental observations we proposed a simple model (Fig. 2g) in which dorsal actin fibers, as in apex 2, keep the nucleus stretched along the fiber direction (through LINC complex connections) while the collapse of the ventral actin fiber network cannot provide structural support for the nucleus in the radial direction of apex 1 and 3. Collapsed ventral actin fibers along the secondary apexes might even contribute to the higher compression observed at their radial positions. Of course, this model neglects other cytoskeleton and other physical components that might have an influence on nuclear strains during the collapse. However, it is important to note that chondrocytes in their native 3D cartilage environment have a sphere-shaped constellation including a spherical actin cytoskeleton that encloses the nucleus. Thus, our model already provides a meaningful explanation for how a spherical actin skeleton prevents local nuclear disturbances during osmotic challenges (e.g. during walking) as observed in the 2D cultures, thereby highlighting the importance of this native 3D spherical confirmation and the need to study chondrocyte mechanosensation in their native environment.

As another application, we conducted intranuclear strain measurements during the spreading of passage 4 chondrocytes (Supplementary note 4, Supplementary Fig. 8). With ongoing cell spreading, an increasingly compressive trend was observed for hydrostatic strain patterns, while shear and deviatoric strains remained mostly unchanged. This was anticipated as cell dehydration can be assumed to be equi-biaxial, and hence changes in volume (as represented by hydrostatic strain) should be dominant.

In conclusion, *deformation microscopy* provides a powerful tool to evaluate intracellular and intranuclear patterns of displacement and strain that relate to distributions of mechanical force through distinct subcellular architectures. This method is neither restricted to the nucleus nor *in vitro* cultures, but can be utilized over a wide spatial range including cells and tissues *in vivo*, as long as image acquisition is sufficient to provide unique features that can be tracked in image sequences. In combination with other quantitative biology techniques, deformation microscopy can enable spatial description of mechanical parameters, i.e. elastography, and be applied to a broad range of mechanobiological applications.

## Materials and Methods

### Validation of deformation microscopy

Images of passage four chondrocytes (P4, exhibiting a fibroblast-like phenotype) were captured by fluorescence confocal microscopy (Nikon A1R) with DNA staining (DRAQ5; Cell Signaling). Z-projections of confocal nucleus images were used in the analysis to enable simulations in two-dimensions, although extension to three-dimensional analysis is possible (*20*). More compact heterochromatin regions were stained brighter compared to euchromatin regions. The euchromatin and heterochromatin regions were segmented using a custom algorithm based on the computation of the second derivative of image intensity values.

Validation of deformation microscopy was determined using known forward simulations of deformation under loading. To account for unknown material properties of nuclear subregions, we assigned different stiffness values to heterochromatin and euchromatin areas defined by segmentation of confocal images. Four different cases were used, representing maximum and minimum stiffness expected in the heterochromatin and euchromatin subregions: nucleus A (H:E = 1:10), nucleus B (H:E = 10:1), nucleus C (H:E = 2:1) and nucleus D (H:E = 1:2). A linear elastic material model was used for the validation and the warping analysis. The nucleus deformation was simulated in ANSYS by applying equal tensile normal force on the nuclear periphery as the boundary condition, so that the magnitude of deformation was similar to that observed in our studies (*20*). The difference between known and computed strain measures at same spatial coordinates were used to validate the *deformation microscopy* technique with a difference of 0 representing perfect registration. The coefficient of determination was calculated between known and quantified strain to further measure the efficacy of hyperelastic warping microscopy.

### Disruption of LINC complexes in cardiomyocytes *in vitro*

Cardiomyocytes were transduced with lentiviral particles harboring a dominant negative sR3-Kash3 construct (Fig 2a). This construct contained the C-terminal end of the nesprin-3 gene (*Syne3*) including the transmembrane (Tm) and the KASH domain. The N-terminus was replaced by a red fluorescence protein (mRuby3) for visual feedback and a signal peptide (SP, from *Tor1a*) for proper membrane integration. This truncated nesprin was designed to integrate into the outer nuclear membrane and bind to SUN proteins via the KASH domain but not to any cytoskeleton components due to the lacking N-terminal.

#### Cardiomyocyte culture

B6.Cg-Tg (HIST1H2BB/EGFP) 1Pa/J mice were obtained from Jackson Laboratory (Bar Harbor, Maine). Animal procedures were performed following Institutional Animal Care & Use Committee approved protocols. CMs were isolated from embryonic mice hearts 16.5 days post conception using cold trypsin digestion (incubation in 0.125% trypsin/EDTA overnight followed by 10 min digestion in residual trypsin under application of 37°C warm medium) and cultured on PDMS substrates.

#### Lentiviral Transduction

Lentiviral particles were produced using Lenti-X Packaging Single Shots (Takara Bio USA, Inc.). For efficient transduction in CM’s, particles where precipitated with 4mM CaCl_2_ in Advanced DMEM/F12 (Thermo Fisher Scientific) without serum and antibiotics for 1h at RT. After 24h of culture, CM’s were incubated in infectious medium for 2h after which cells were switched to complete CM medium again. After 4 days of transduction, nuclear image stacks of CM’s with strong marker expression or non-transduced control cells were acquired and analyzed via deformation microscopy as described before. Target sequences for shRNA constructs can be obtained from Supplemental Table 1.

#### Nesprin Immunostaining

Cells were fixed in 4% PFA for 10min, blocked with 10%NGS, 1% BSA in 0.1% PBT (0.1% Tween-20 in PBS) for 60min. Primary incubation was performed at 4°C overnight in 0.1% PBT containing 1% BSA. Primary antibodies: nesprin-1 (Abcam, ab24742, 1:500); nesprin-2 (Santa Cruz, sc-99066, 1:100). Secondary incubation was performed in primary incubation buffer for 40min RT at a dilution 1:500. Images were acquired on a Nikon A1R confocal microscope using a 60x oil immersion objective.

#### Gene Expression Analysis

Total RNA was extracted from CMs 4 days after transduction using AurumTM Total RNA Mini Kit, was reverse transcribed into cDNA via iScriptTM Reverse Transcription Supermix and real-time quantitative PCR was performed with SsoAdvancedTM Universal SYBR® Green Supermix in a CFX96 Touch™ thermocycler (all kits and devices from Bio-Rad Laboratories) using 10 ng of cDNA as input. Primers were custom designed using NCBI primer blast, cross-confirmed in Ensembl gene database and synthesized by IdtDNA. All primers span at least one exon-exon junction. Relative expression change was calculated using the ΔΔCt method. All data was normalized to the reference gene Gapdh. Primer sequences are listed in Supplemental Table 1.

### Quantification of skeletal muscle nuclear deformation *in vivo* in Lmna knockout mouse

All animal experiments were performed under Institutional Animal Care and Use Committee (IACUC) approved protocols. B6129S1(Cg)-*Lmna*^*tm1Stw*^/BkknJ mice were purchased from Jackson Laboratory (Bar Harbor, Maine). The targeted allele does not express both full-length transcripts or stable lamin A/C protein. Mice heterozygotes for this lamin A/C mutation (*Lmna*^*+/-*^) are viable and fertile. Heterozygotes were bred to obtain the *Lmna*^*+/+*^, *Lmna*^*+/-*^, *Lmna*^*-/-*^ mice. Homozygotes (*Lmna*^*-/-*^ mice) exhibit severely retarded postnatal growth beginning as early as 2 weeks of age and abnormal movement/gait by 3-4 weeks of age. These mice are a model for the autosomal variant of muscular dystrophy and are useful in studying the role of lamin A/C.

Skeletal muscle of the *Lmna*^*-/-*^ mice show compromised nuclear phenotype. We applied a neuromuscular stimulation *in vivo* to quantify the nuclear strain in skeletal muscle as described previously (*36*). Briefly, anesthetized animals were kept in supine position and their gastrocnemius was exposed. The nuclei were stained by NucBlue Live ReadyProbes Reagent (Thermo Fisher Scientific Inc). On the stimulation of the deep fibular nerve, the gastrocnemius activated and deformed. Nuclei in the medial gastrocnemius were imaged before and after deformation (activation) using an inverted confocal microscope (Nikon Eclipse Ti A1R) at 40× magnification, and were quantified further using deformation microscopy.

### Cardiomyocyte (CM) culture and beating experiment

#### Substrate fabrication

To mimic the normal and pathological conditions, two different PDMS formulations (Dow Corning) were used: Sylgard®527 ratio 1:1 (*E* = 11.7 ± 5.4 kPa) and Sylgard®184 ratio 1:10 (*E* = 434.3 ± 54.4 kPa). PDMS Stiffness was determined via AFM using a spherical borosilicate glass tip (diameter = 10 µm, stiffness = 0.85 N/m), and Young’s modulus *E* was calculated using a Hertz contact model. To enable live imaging at high magnification using a 100× objective, thin (∼80 µm) PDMS films were deposited on glass slides. PDMS substrates were degassed under vacuum for 30 min, cured for 2h at 100°C, and mounted on custom made cell culture dishes. PDMS was then ozone-treated and coated with Matrigel (Geltrex®, ThermoFisher) for 1h at 37°C to allow for cell attachment.

#### Imaging and Data analysis

After 4 days of CM culture on PDMS, images stacks were captured using an inverted epi-fluorescence microscope (Nikon Ti-Eclipse) with a 100× objective and an EMCCD camera (Andor). To visualize the entire contraction cycle of CMs, images were captured at 6.4 fps over a period of 10 s, during which photobleaching was negligible. For each nucleus, one contraction cycle was selected from the image stack to perform nuclear deformation analysis. Images of the nuclei in a post-systolic resting state were selected as undeformed reference image. Statistical measures over multiple nuclei (*n*=5) were determined using JMP Pro12 software (SAS Institute).

### Hyperosmotic loading of chondrocytes, imaging and analysis

#### Chondrocyte culture

Primary bovine chondrocytes were harvested from the load-bearing region of the medial condyle from young bovines (*37, 38*). Harvested cells were cultured in DMEM-F12 with 10% FBS and passaged at around 80% confluency.

#### Substrate fabrication

Polydimethylsiloxane (PDMS; Sylgard®184; Dow Corning) were mixed with 0.5 µm red fluorescent beads (ThermoFisher) and sandwiched between a petri dish and a glass slide. Substrates were degassed for 20 min, cured at 70°C overnight, and coated with 1µg/cm^2^ fibronectin (Life Technologies) for 1 hour.

#### Imaging setup

Cells were plated at low density (i.e. maximum of ∼5 to 9 cells per 106×106 µm^2^) on substrates for 2 hours prior to hyperosmotic loading experiments, and cell nuclei were labeled with Hoechst 33342 (ThermoFisher). Imaging was performed using a two-photon confocal microscope (Olympus) with a tunable Mai Tai pulse laser (excitation wavelength 740 nm), and a 60× water immersion objective. The detector wavelength was set to 430-500 nm (green) and 540-600 nm (red) to image cell nuclei and the fluorescent beads, respectively. The green channel also detected the cell shape through NADH autofluorescence.

#### Hyperosmotic loading of chondrocytes

After placing the substrate on confocal microscope, images of the nucleus and the substrate were captured. The culture media was removed, and prior to cell dehydration, saline solution was added (dropped gently) on the cell to increase the osmolality from 320 mOsm to 500 mOsm instantly, thus creating a hyperosmotic shock. Subsequently, 10 images of the same nucleus and the substrate were captured one minute apart. Finally, the cell was detached from the substrate using TrypLE Express (Life Technologies) and another set of images was captured that showed only the beads in the substrate. A control experiment was conducted using the regular culture medium for the same time span as the hyperosmotic loading experiment.

#### Data analysis for traction force, stretching force on cell, and nuclear strain

The image sequence of the nuclei obtained from hyperosmotic loading experiment was further analyzed to compute nuclear strain. From the substrate images, traction force and substrate stress relaxation force on cell was computed. Briefly, nuclear deformation was computed for each time point with respect to the first nuclear image (before application of osmotic loading) as the template. Traction forces at each time point were computed via the ImageJ-based PIV and traction force calculation plugin (*27*) with reference to the last image after trypsin-mediated cell detachment. Substrate stress relaxation maps were computed by subtracting the traction force at the initial time point (0 min) from traction force maps at each time point (*39*). Young’s modulus of the PDMS was estimated to be *E* = 1.758 MPa, and Poisson’s ratio *v* = 0.49 based on existing literature (*40*). A simple measure of bulk nuclear strain was obtained by fitting an ellipse to the nucleus and computing the change in major axis length.

### Statistics

One-way Analysis of Variance (ANOVA), followed by post hoc Tukey’s test was used to find any statistically significant difference between the groups. The coefficient of regression (*R*^*2*^) was calculated using linear regression.

## General

We thank Jessica Kelly and Adrienne Kathleen Scott for technical assistance.

## Author contributions

All authors conceived of the study, designed the experiments, and reviewed the manuscript. SG, BS, RW, and JH performed the experiments and carried out the data analysis. SG, BS, and CPN wrote the manuscript.

## Supplementary Materials

### Supplementary Note 1: Deformable image registration and hyperelastic warping

Consider a space variable *X*, a displacement map *u*(*X*), and a deformation map*φ*(*X*) defined as

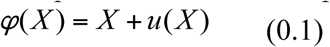

Deformable image registration methods aim to find the deformation map *φ*(*X*) which represents minimization of the energy functional, *E*(*X, φ*), that represents the difference in the strain energy within the deforming material, *W* (*X, φ*), and an image based energy term, *U* (*X, φ*), which is a pointwise comparison of the undeformed (template) image *T* and the deformed (target) image *S*. This can be mathematically expressed by the following.

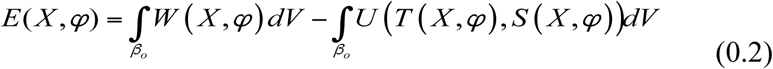

where the first term on right side is the energy function of the deformation map and the second term is the energy function comprising image comparison. The terms are evaluated at every differential image location *dV*, and is integrated over the whole image space *β*_*o*_. For hyperelastic warping, the first term *W* is the standard energy density function modeled as dependent on the material constitutive model, material properties, and right Cauchy-Green tensor *C*, defined as *C* = *F^T^ F*, where *F* is the deformation gradient tensor,

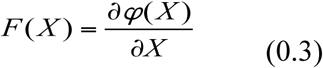

The second term *U* uses a Gaussian tensor model to describe the image energy density functional.

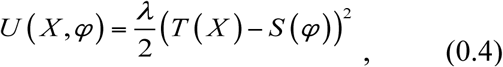

where *T* is the scalar intensity field of the template and hence a function of *X* only. *S* is the scalar intensity of the target, and hence a function of *φ*. *λ* is a penalty parameter that enforces the alignment of the template with the target. Details of hyperelastic warping can found elsewhere^1^.

### Supplementary Note 2: Details of deformation microscopy

#### 2.1 Parameter sweep

The goal of deformation microscopy is to reach a pixel intensity matching solution based on the warped template and target images. The iterative hyperelastic warping algorithm is run over a range of two parameters: the material stiffness (range: 100-10000 Pa, with a minimum step size of 100) and the penalty (range: 1-3.5, with a step size of 0.5). For each set of parameters, the root mean square error (RMSE) in intensity difference based on warped template and target is quantified. The RMSE value was minimized over the parameter sweep and thus the best possible set of parameters was obtained. A custom made MATLB code was written to automate and connect the software for mesh generation (TrueGrid), deformable image registration (nike3D) and postprocessing (WarpLab and LSPrePost). By using parallel solver for the different penalty parameters, the iterative procedure can be performed in tens of minutes for one nucleus. Supplementary Fig. 1 explains the iterative procedure and Supplementary Fig. 2c shows the results of a sequential iterative process to reach the final deformation map.

#### 2.2 Sequential spatial filtering

Local minima in the registration process can lead to premature registration for some local image features thus ending up with inaccurate and incomplete registration. To overcome this, sequential low pass filtering is used. First, the larger image features, e.g. object boundaries and coarse texture details, are registered (global registration) followed by registration of finer details (local registration) by gradually reducing the extent of the spatial filter. The spatial filter is applied by convolution of the image with a kernel *ĸ* (*X*). For the image field *T*,

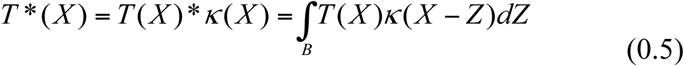

where *Z* is the frequency domain variable of *X* and *T* *(*X*) is the spatial intensity data in the filtered image in the iteratively deformed domain *B*. In this particular study, a 3D Gaussian kernel has been used.

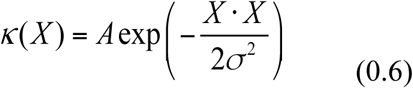

where *σ*^2^ constant. is the spatial variance used to control the extent of blurring and *A* is a normalizing

#### 2.3 Underregistration and overregistration

During the deformable registration process, the deformed template may be underregistered at certain parameter values, despite the blurring applied to that that particular parameter. This leads to a false minimum total energy *E* and the deformation values we get is not optimal. On other hand if in equation 1.2, the *U* term is bigger than the *W* term, the minimized energy becomes negative in some spaces and a false deformation is reported termed as ‘overregistration’. This is demonstrated in Supplementary Fig. 2, for the two parameters stiffness (S2.a) and penalty (S2.b).

### Supplementary Note 3: Strain measures to quantify deformation

Deformable microscopy directly yields displacement measures, which leads to fundamental strain measures. For two-dimensional (2D) analysis, as performed in this work using the z projection of the images, the relevant strain measures computed are: *E*_*xx*_, *E*_*yy*_ and *E*_*xy*_.

The hydrostatic strain can be derived from those quantities as:

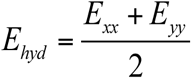

The first and second deviatoric strains are computed as:

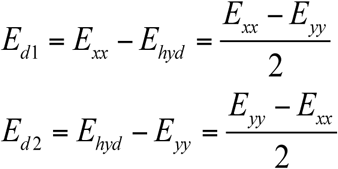

Therefore, *E*_*d*1_ = – *E*_*d*2_ for 2D measurement. Unless otherwise mentioned all quantifications of deviatoric strain in this study are the second deviatoric strain. Note that this relation does not hold true for three-dimensional (3D) measurements. For this work, all nucleus deformation experiments are performed on cells plated on 2D surface, therefore the z directional strain has minimum effect on the two dimensional strain measurement. However, these methods can be applied and extended to 3D strain measurement as reported^2^.

### Supplementary Note 4: Nuclear deformation during cell spreading

Passage 4 chondrocytes were derived as described in the chondrocyte hyperosmotic loading section. DNA of passage four bovine chondrocytes were stained in suspension with DRAQ5 (Cell Signaling). Chondrocytes were then plated on plastic dishes maintained at 37°C, while two-photon microscopy (Olympus with a tunable Mai-Tai laser set at 540-600 nm) acquired time-lapse images of the cells (transmitted light) and nuclei (fluorescence). See Figure S7.a for details of the cell spreading. Intuitively, this simple cell spreading assay predicts a uniform strain pattern in the nucleus, as there is no directional loading applied on the cell. Bulk mechanical behavior of cell computed by cell aspect ratio and cell nuclear area follows this predictable pattern. Nucleus aspect ratio was near 1 (S7.b), suggesting a biaxial expansion and contraction during the time-course change of the nucleus area. However, the internal strain fields were locally complex, with spatiotemporal variations in tensile and compressive strain magnitudes, thus leading to complex hydrostatic and deviatoric strain (S7.c). From the spreading to anchoring phase, at some time points, high average tension (S7.d) in cell was found, suggesting cells attach on the substrate by creating short protrusions. These results warrant further investigation of local nuclear behavior during cell spreading.

**Figure S1:**
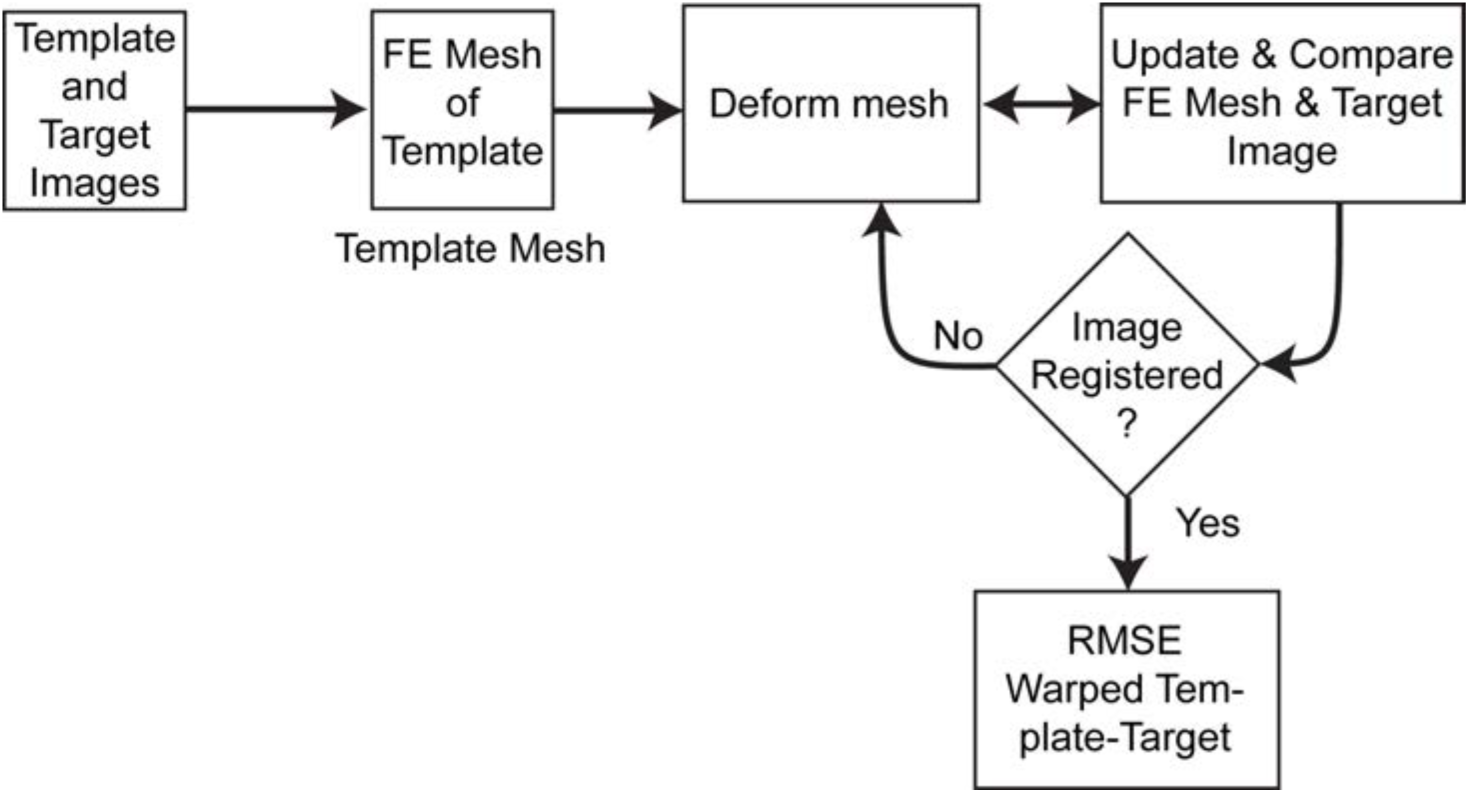
Flowchart explaining iterative hyperelastic warping based deformable image registration. The procedure is repeated with parameter (stiffness and penalty) sweep.

**Figure S2:**
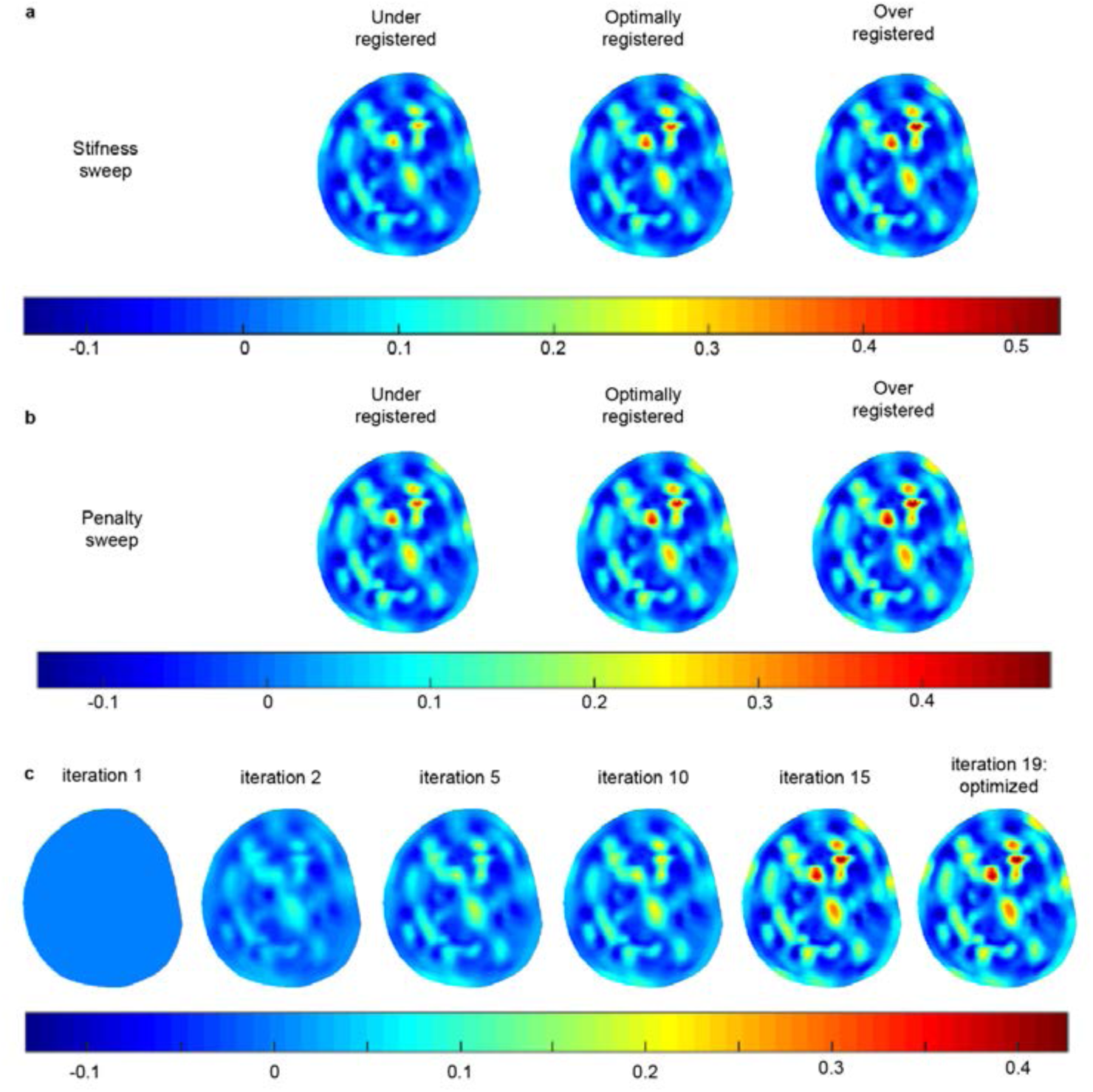
Registration process during iterative hyperelastic warping. Demonstration of underregistration and overregistration for the two parameters : stiffness **(a)** and penalty **(b)** through hydrostatic strain map. **(c)** Shows how an optimal solution is reached by the iterative procedure after best registration is achieved.

**Figure S3:**
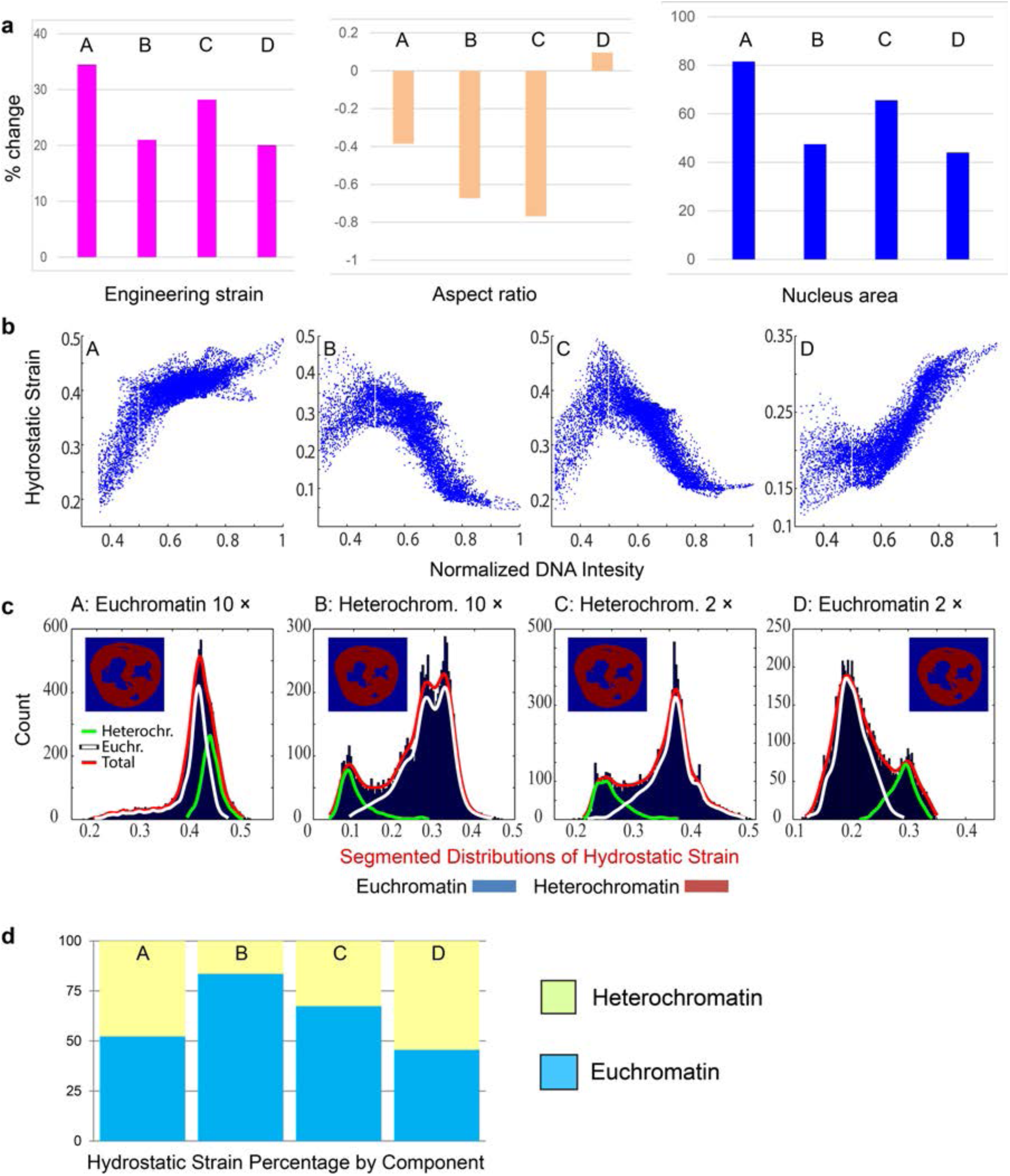
Effect of intranuclear stiffness on strain measurement. **(a)** Percent strain of the four nuclei A, B, C and D as measured by bulk measurements (i.e. engineering strain based on major axis, aspect ratio and nucleus area). **(b)** Hydrostatic strain vs. normalized chromatin intensity. For nuclei A and D, where the euchromatin region is stiffer, similar strain vs intensity trends were observed, with opposite trends found for nuclei B and C. **(c)** Segmentation of the nucleus images into euchromatin and heterochromatin regions show that the total strain distribution (blue histogram, highlighted with a red line) is a composite of the two strain distribution representing heterochromatin (green line) and euchromatin (white line). **(d)** Percentage of total hydrostatic strain distributed between the hetero- and euchromatin components.

**Figure S4:**
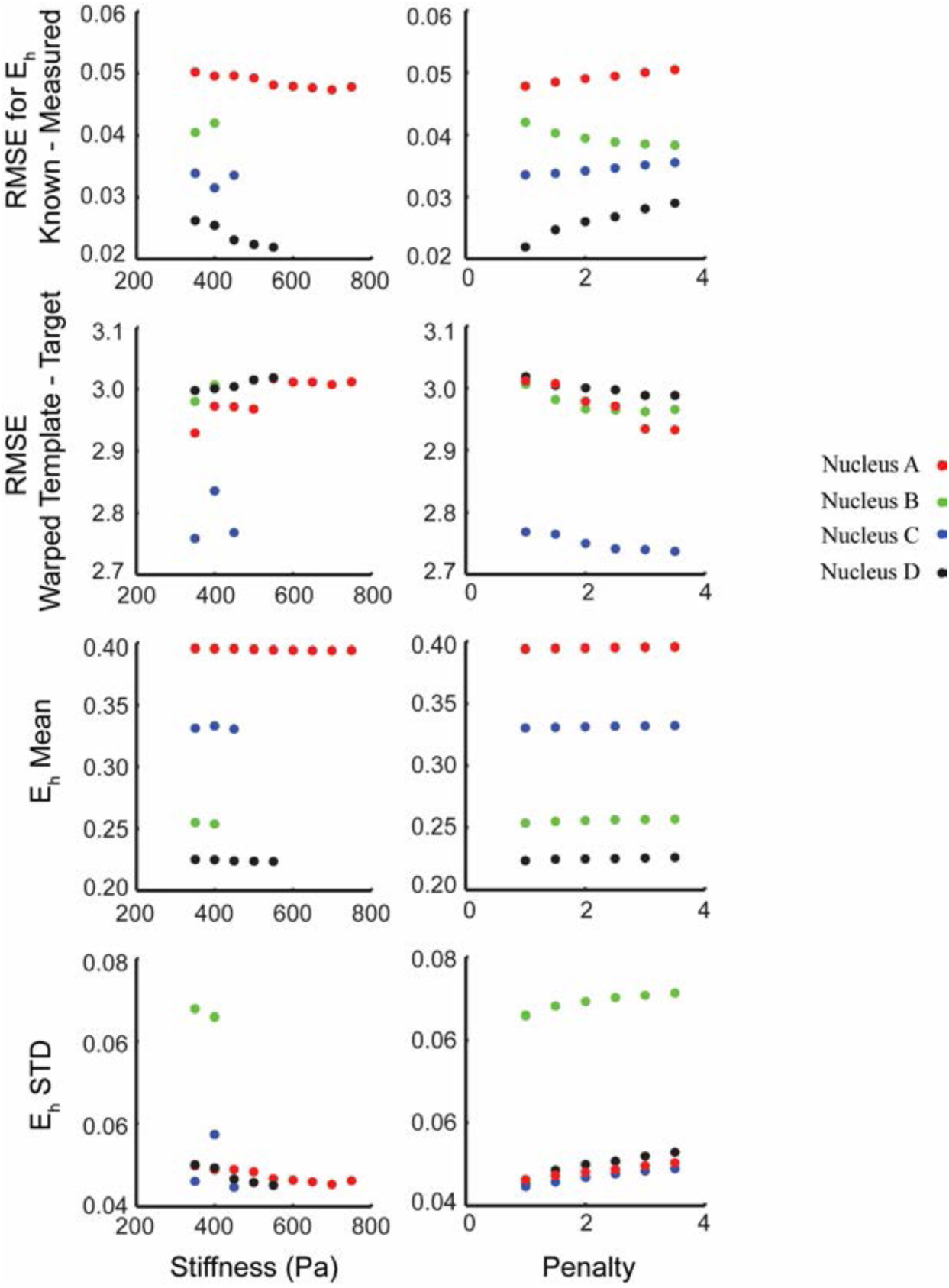
Efficacy of parameter sweep in deformation microscopy. To quantify the efficacy of our new technique, the root mean square error (RMSE) was computed comparing known forward simulation generated hydrostatic strain and the hydrostatic strain measured by deformation microscopy. Root mean square error (RMSE) was also computed for image intensity of the warped template and the target. The mean and standard deviation of the hydrostatic strain value was also computed for the nucleus. The results are shown for stiffness and penalty parameter sweep for all four nuclei. The RMSE reaches a minimum with increasing stiffness and penalty, thus reaching the best possible solution. The average of hydrostatic strain is insensitive to both the change in stiffness and penalty. The standard deviation in hydrostatic strain increases with increasing penalty but decreases with increasing stiffness.

**Figure S5:**
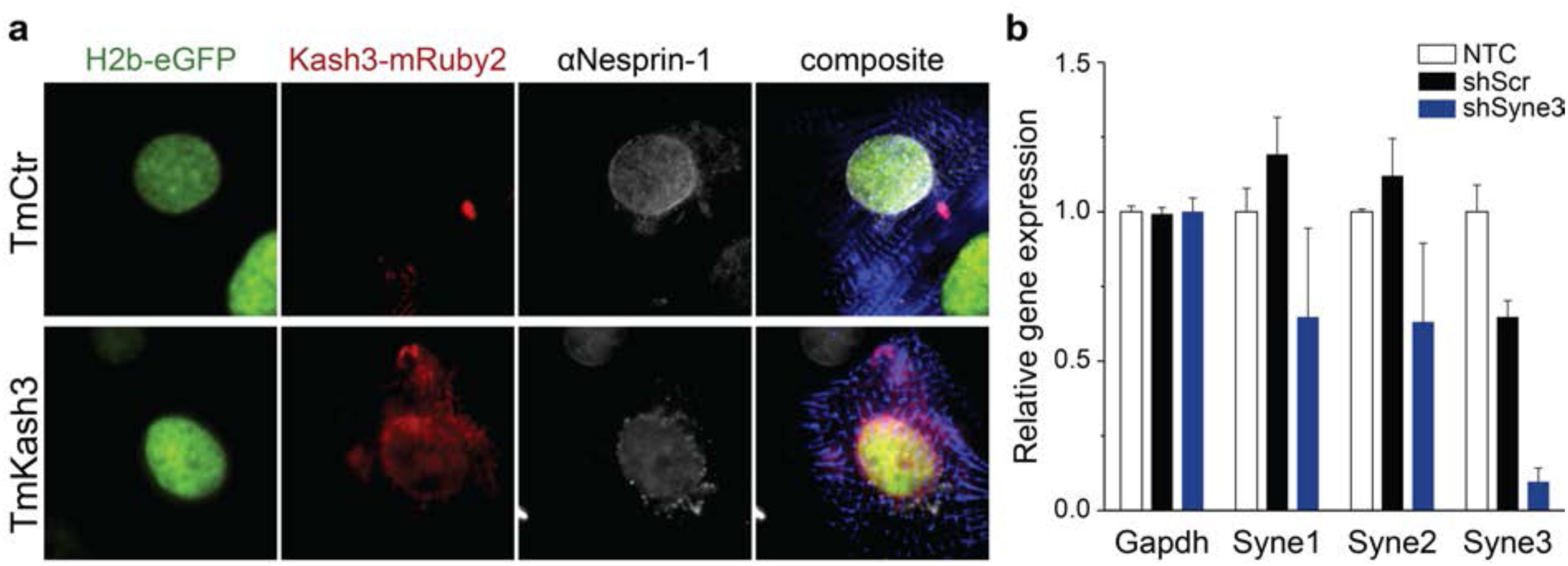
Validation of LINC complex disruption via TmKash3 or shSyne3 expression in cardiomyocytes. **A)** Immunofluorescence staining of Nesprin-1 in cardiomyocytes transduced with either TmCtr or TmKash3. The truncated nesprin (Kash3-mRuby2) expression construct integrated properly around the nucleus and disrupted the localization of Nesprin-1 to the nuclear membrane in turn. B) PCR gene expression analysis of Nesprins 1 to 3 between cardiomyocytes transduced with an shRNA cassette against a random scrambled sequence (shScr) or against Syne3 (shSyne3) or non-transduced control cells (NTC); error bars=STD, n=4.

**Figure S6:**
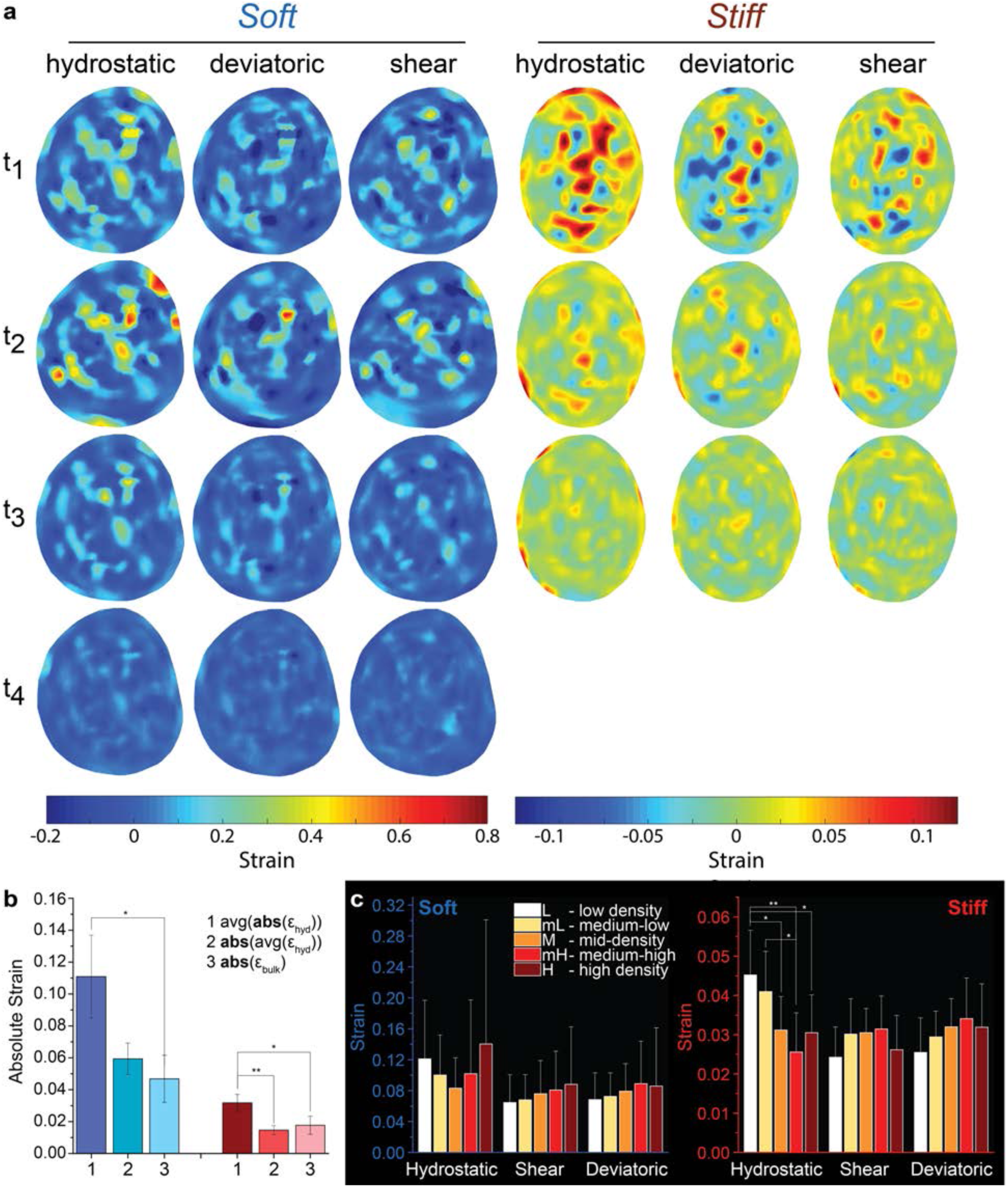
Comparison of strain map obtained using deformation microscopy and bulk strain measurements for cardiomyocyte nuclei. Deformation microscopy analysis of cardiomyocyte nuclei plated on either soft (~15 kPa) or stiff (~150 kPa) PDMS substrates. **(a)** Spatial strain maps of cardiomyocyte nuclei engineering strain based on major axis, **(b)** Comparison of deformation microcopy and bulk strain analysis. **(c)** Average strain in each chromatin density segment for nuclei in CMs platted on soft or stiff substrates; (b, c) ** p<0.01, * p<0.05, otherwise non-significant, n=5.

**Figure S7:**
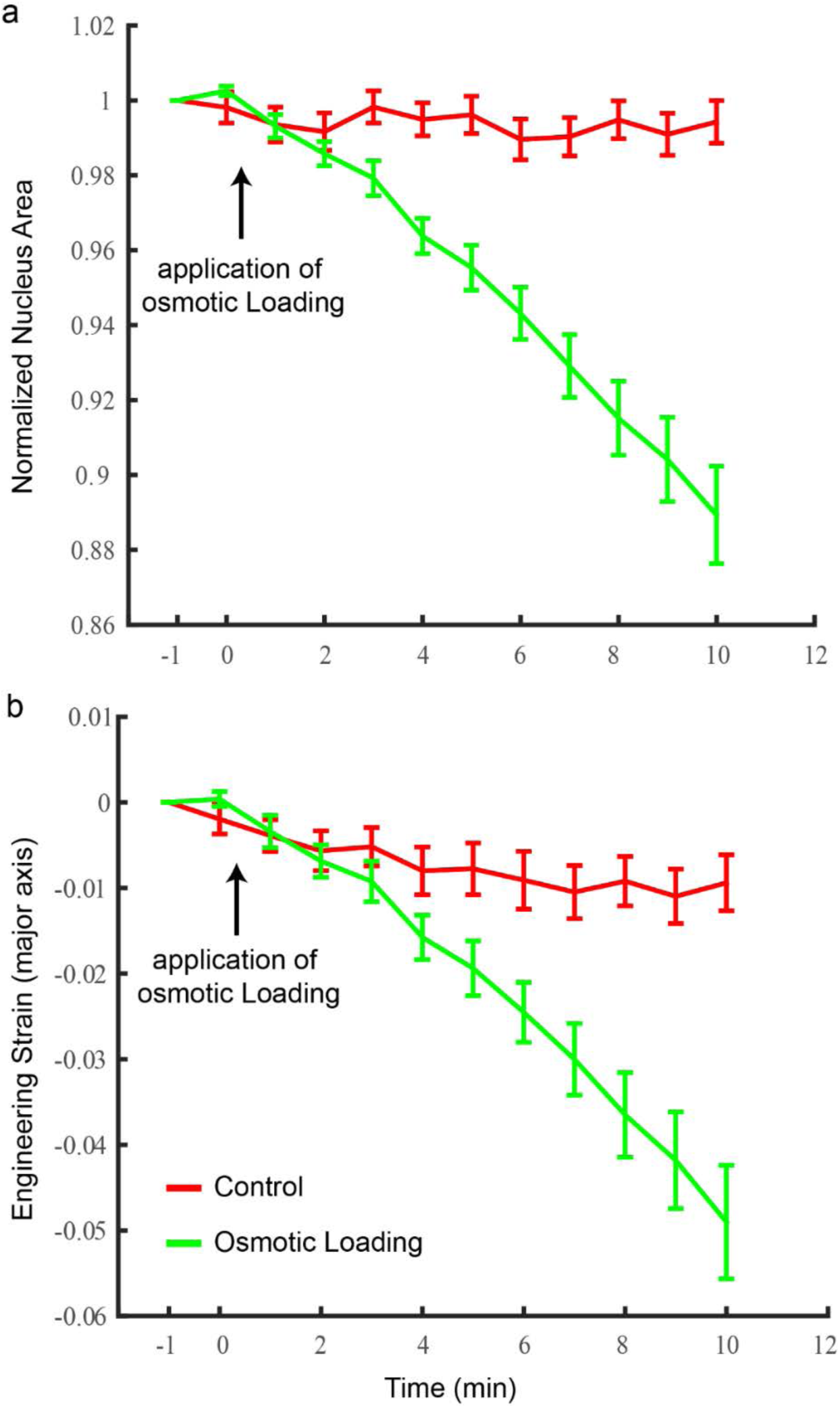
Bulk mechanical strain measurement during chondrocyte osmotic loading. Bulk mechanical strain measurements: **(a)** normalized area and **(b)** engineering strain during osmotic loading of passage 4 chondrocytes over time. As water diffuses out of the cell and nucleus both measures show overall increase in negative compressive strain with time (green). Control cells in medium shows no significant change in those quantities (red); lines and bars represent means and standard deviation, osmotic loading: n=27, control: n=32.

### Supplementary Figure 8

**Figure S8:**
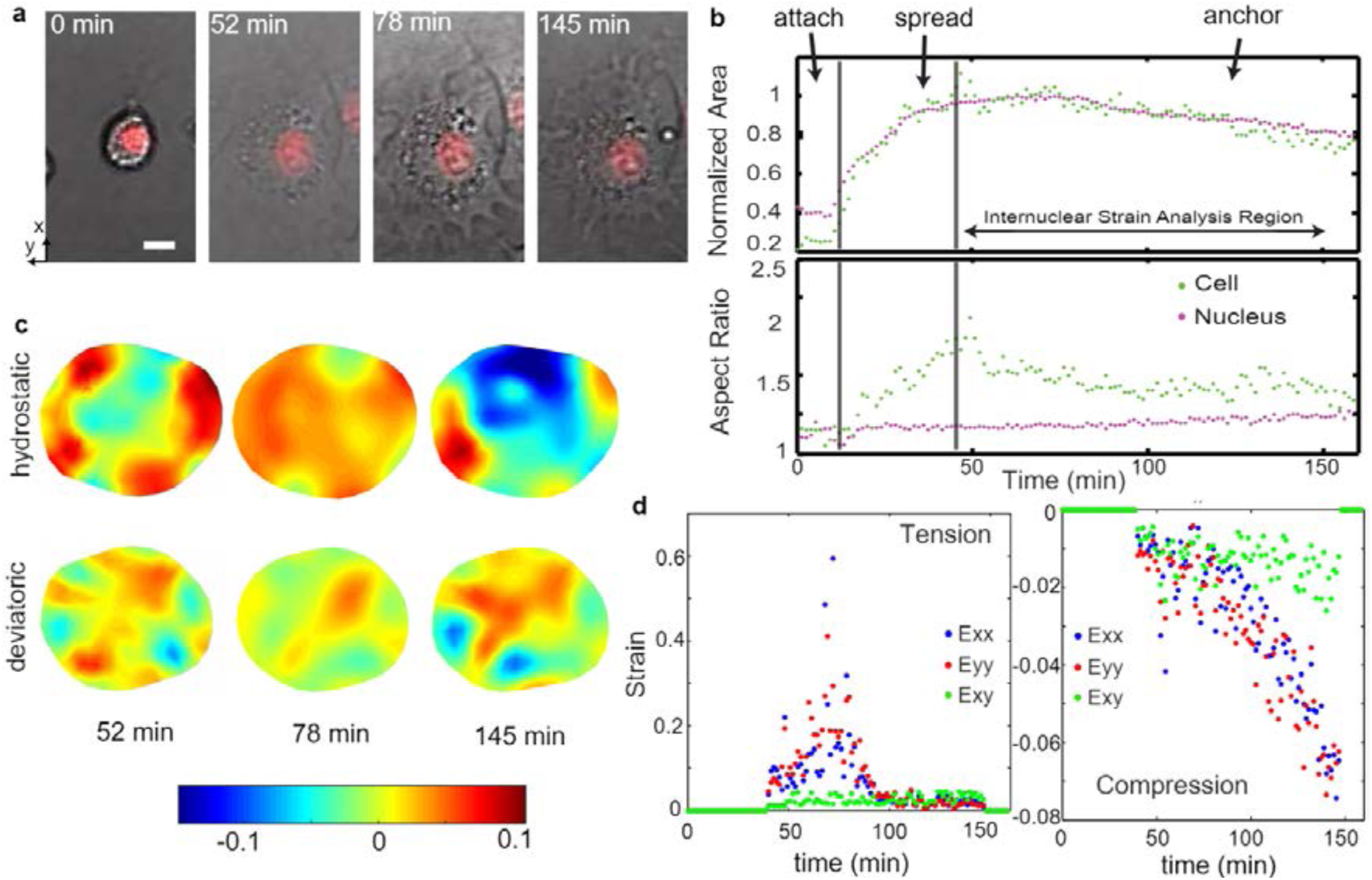
Nuclear deformation during cell spreading. Deformation microscopy and bulk strain analysis of a P4 chondrocyte nucleus during cell spreading. **(a)** Brightfield microscopy of cell (transmitted) nucleus (red) during cell spreading. **(b)** Area and aspect ratio change of cell and nucleus indicates different phases during cell spreading. Area and aspect ratio were computed by masking cell and nucleus images in ImageJ. **(c)** Intranuclear strain maps computed via deformation microscopy using the start of the anchoring phase (49 min) as undeformed reference. Average tensile and compressive strains over time. Tensile strain showed a temporal spurt from spreading to anchoring phase while compressive strain steadily increased with time.

**Supplemental Table 1.**
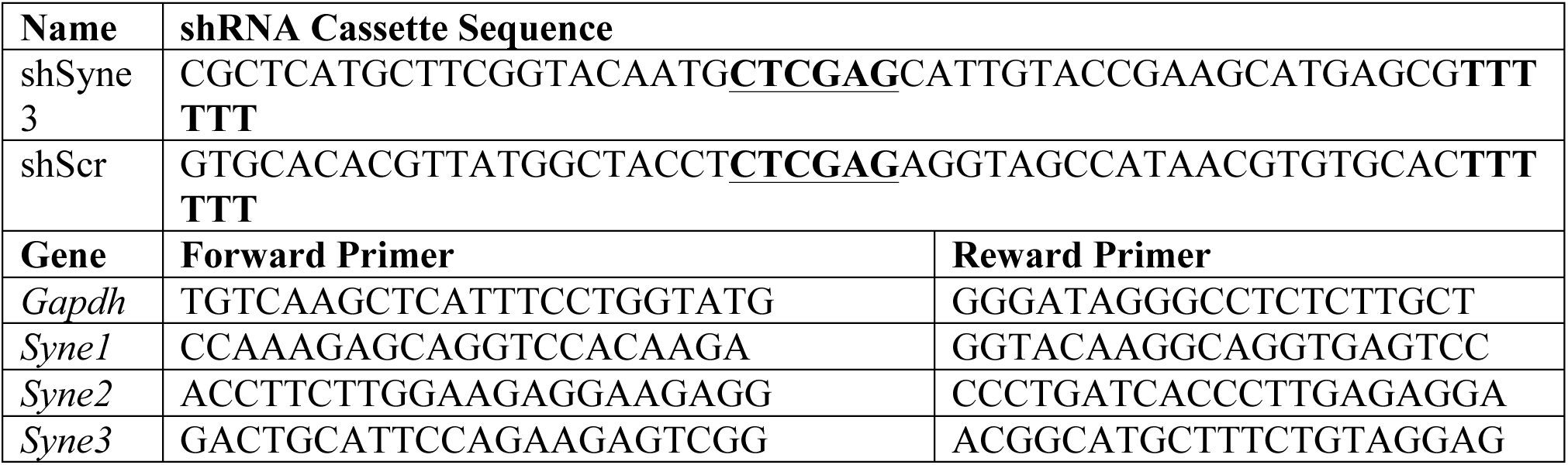
Primer and shRNA hairpin sequences. Top: Hairpins with target sequence against nesprin-3 or scrambled control sequence. Hairpin is underlined and bold. Bottom: Primer sequences for qPCR validation of shRNA-mediated nesprin knock-down.

**Supplementary video 1 Time-lapse microscopy of hyper-osmotically dehydrating chondrocyte on a substrate embedded with beads.** Nucleus is shown in green, beads are red and the cell contour can be visualized in bright field, at 7 min timepoint parts of the cell contour marked with pink arrows. The microscopic image stack was analyzed to quantify the substrate stress relaxation using traction force microscopy and nuclear strain using deformation microscopy as explained in the main text, Figure 2.

**Supplementary video 2 Time lapse microscopy of cardiomyocyte beating on soft vs stiff substrates.** The nucleus deforms more on soft substrate and less on stiff substrate, the extent of mechanical strain is quantified using deformation microscopy. See main text, Figure 3. Scale bar: 10 µm.

## References

1. T. Iskratsch, H. Wolfenson, M. P. Sheetz, Appreciating force and shape—the rise of mechanotransduction in cell biology. Nat. Rev. Mol. Cell Biol. 15, 825–33 (2014).

2. N. Wang, J. D. Tytell, D. E. Ingber, Mechanotransduction at a distance: mechanically coupling the extracellular matrix with the nucleus. Nat. Rev. Mol. Cell Biol. 10, 75–82 (2009).

3. C. Guilluy et al., Isolated nuclei adapt to force and reveal a mechanotransduction pathway in the nucleus. Nat. Cell Biol. 16, 376–381 (2014).

4. C. M. Denais et al., Nuclear envelope rupture and repair during cancer cell migration. Science (80-.). 352, 353–358 (2016).

5. A. Tajik et al., Transcription upregulation via force-induced direct stretching of chromatin. Nat. Mater.(2016), doi:10.1038/nmat4729.

6. M. L. Lombardi et al., The interaction between nesprins and sun proteins at the nuclear envelope is critical for force transmission between the nucleus and cytoskeleton. J. Biol. Chem. 286, 26743–53 (2011).

7. S. G. Alam et al., The mammalian LINC complex regulates genome transcriptional responses to substrate rigidity. Sci. Rep. 6, 38063 (2016).

8. T. P. Driscoll, B. D. Cosgrove, S.-J. Heo, Z. E. Shurden, R. L. Mauck, Cytoskeletal to Nuclear Strain Transfer Regulates YAP Signaling in Mesenchymal Stem Cells. Biophys. J. 108, 2783–2793 (2015).

9. I. Banerjee et al., Targeted ablation of nesprin 1 and nesprin 2 from murine myocardium results in cardiomyopathy, altered nuclear morphology and inhibition of the biomechanical gene response. PLoS Genet. 10, e1004114 (2014).

10. J. Swift et al., The nuclear lamina is mechano-responsive to ECM elasticity in mature tissue. J. Cell Sci. 127, 3005–15 (2014).

11. K. V. Iyer, S. Pulford, A. Mogilner, G. V. Shivashankar, Mechanical activation of cells induces chromatin remodeling preceding MKL nuclear transport. Biophys. J. 103, 1416–1428 (2012).

12. S. J. Heo et al., Biophysical regulation of chromatin architecture instills a mechanical memory in mesenchymal stem cells. Sci. Rep. 5, 16895 (2015).

13. Y. Jamali, R. Karimi, M. R. K. Mofrad, Biophysical Coarse-Grained Modeling Provides Insights into Transport through the Nuclear Pore Complex, doi:10.1016/j.bpj.2011.01.061.

14. K. N. Dahl, A. J. Engler, J. D. Pajerowski, D. E. Discher, Power-Law Rheology of Isolated Nuclei with Deformation Mapping of Nuclear Substructures. Biophys. J. 89, 2855–2864 (2005).

15. M. Zwerger et al., Myopathic lamin mutations impair nuclear stability in cells and tissue and disrupt nucleo-cytoskeletal coupling. 22, 2335–2349 (2017).

16. P.-H. Wu et al., A comparison of methods to assess cell mechanical properties. Nat. Methods (2018), doi:10.1038/s41592-018-0015-1.

17. M. M. Knight et al., Cell and nucleus deformation in compressed chondrocyte-alginate constructs: Temporal changes and calculation of cell modulus. Biochim. Biophys. Acta-Gen. Subj. 1570, 1–8 (2002).

18. C. L. Gilchrist, S. W. Witvoet-Braam, F. Guilak, L. A. Setton, Measurement of intracellular strain on deformable substrates with texture correlation. J. Biomech. 40, 786–794 (2007).

19. S. Talwar, A. Kumar, M. Rao, G. I. Menon, G. V. Shivashankar, Correlated spatio-temporal fluctuations in chromatin compaction states characterize stem cells. Biophys. J. 104, 553–564 (2013).

20. J. T. Henderson, G. Shannon, A. I. Veress, C. P. Neu, Direct measurement of intranuclear strain distributions and RNA synthesis in single cells embedded within native tissue. Biophys. J. 105, 2252–61 (2013).

21. K. K. Brock, M. B. Sharpe, L. a Dawson, S. M. Kim, D. a Jaffray, Accuracy of finite element model-based multi-organ deformable image registration. Med. Phys. 32, 1647–1659 (2005).

22. A. (U of U. Veress, N. (U of U. Phatak, J. (U of U. Weiss, Deformable Image Registration with Hyperelastic Warping. Handb. Med. Image Anal. Segmentation Regist. Model., 487–534 (2004).

23. I. Sarantitis, P. Papanastasopoulos, M. Manousi, N. G. Baikoussis, E. Apostolakis, The cytoskeleton of the cardiac muscle cell. Hellenic J. Cardiol. 53, 367–79 (2012).

24. T. Lanzicher et al., AFM single-cell force spectroscopy links altered nuclear and cytoskeletal mechanics to defective cell adhesion in cardiac myocytes with a nuclear lamin mutation AFM single-cell force spectroscopy links altered nuclear and cytoskeletal mechanics to defecti. 1034 (2015), doi:10.1080/19491034.2015.1084453.

25. A. J. Engler et al., Embryonic cardiomyocytes beat best on a matrix with heart-like elasticity: scar-like rigidity inhibits beating. J. Cell Sci. 121, 3794–802 (2008).

26. M. J. Peffers, P. I. Milner, S. R. Tew, P. D. Clegg, Regulation of SOX9 in normal and osteoarthritic equine articular chondrocytes by hyperosmotic loading. Osteoarthr. Cartil. 18, 1502–1508 (2010).

27. J.-L. Martiel et al., Measurement of cell traction forces with ImageJ. Methods Cell Biol. 125, 269–87 (2015).

28. J. Califano, C. Reinhart-King, Substrate stiffness and cell area predict cellular traction stresses in single cells and cells in contact. Cell. Mol. Bioeng. 3, 68–75 (2011).

29. F. Guilak, J. R. Tedrow, R. Burgkart, Viscoelastic Properties of the Cell Nucleus. 786, 781–786 (2000).

30. P. Isermann, J. Lammerding, Nuclear mechanics and mechanotransduction in health and disease. Curr. Biol. 23 (2014), doi:10.1016/j.cub.2013.11.009.Nuclear.

31. J. T. Morgan et al., Nesprin-3 regulates endothelial cell morphology, perinuclear cytoskeletal architecture, and flow-induced polarization (2011), doi:10.1091/mbc.E11-04-0287.

32. R. J. Petrie, H. Koo, K. M. Yamada, Generation of compartmentalized pressure by a nuclear piston governs cell motility in a 3D matrix. 345, 1062–1066 (2014).

33. J. Lammerding et al., Lamin A / C deficiency causes defective nuclear mechanics and mechanotransduction. 113 (2004), doi:10.1172/JCI200419670.Introduction.

34. M. Lawrence, S. Daujat, R. Schneider, Lateral Thinking: How Histone Modifications Regulate Gene Expression. Trends Genet. 32, 42–56 (2016).

35. E. J. Cox, S. A. Marsh, A systematic review of fetal genes as biomarkers of cardiac hypertrophy in rodent models of diabetes. PLoS One. 9, e92903 (2014).

36. S. Ghosh et al., In Vivo Multiscale and Spatially-Dependent Biomechanics Reveals Differential Strain Transfer Hierarchy in Skeletal Muscle, doi:10.1021/acsbiomaterials.6b00772.

37. C. P. Neu, A. H. Reddi, K. Komvopoulos, T. M. Schmid, P. E. Di Cesare, Increased friction coefficient and superficial zone protein expression in patients with advanced osteoarthritis. Arthritis Rheum. 62, 2680–2687 (2010).

38. C. P. Neu, A. Khalafi, K. Komvopoulos, T. M. Schmid, A. H. Reddi, Mechanotransduction of bovine articular cartilage superficial zone protein by transforming growth factor?? signaling. Arthritis Rheum. 56, 3706–3714 (2007).

39. J. M. Mann, R. H. W. Lam, S. Weng, Y. Sun, J. Fu, A silicone-based stretchable micropost array membrane for monitoring live-cell subcellular cytoskeletal response. Lab Chip. 12, 731–40 (2012).

40. I. D. Johnston, D. K. McCluskey, C. K. L. Tan, M. C. Tracey, Mechanical characterization of bulk Sylgard 184 for microfluidics and microengineering. J. Micromechanics Microengineering. 24, 35017 (2014).

## Supplementary References

1. Veress, A., Phatak, M. & Weiss, J. in The Handbook of Medical Image Analysis: Segmentation and Registration Models 587–534 (2004).

2. Henderson, J. T., Shannon, G., Veress, A. I. & Neu, C. P. Direct measurement of intranuclear strain distributions and RNA synthesis in single cells embedded within native tissue. Biopys. J. 105, 2252–2261 (2013).

